# Short-term forecasts of *Aedes aegypti* relative abundance to enhance mosquito control situational awareness

**DOI:** 10.64898/2026.07.03.736030

**Authors:** Utkarsh Bhosekar, Paulo C. Ventura, Megan D Hill, Allisandra G. Kummer, Shreeya Mhade, Jagadeesh Chitturi, Chalmers Vasquez, John-Paul Mutebi, John Townsend, Maria Litvinova, Andre B.B. Wilke, Marco Ajelli

## Abstract

Conventional mosquito surveillance typically relies on contemporaneous data, making it challenging to anticipate future vector surges. To support proactive vector management, this study evaluates a multi-model forecasting framework designed to generate probabilistic 1-to 4-week-ahead forecasts of *Aedes aegypti* relative abundance per trap night. The framework was validated using multi-year surveillance data across four US jurisdictions spanning varied environments (from subtropical to temperate and arid). We found that an ensemble approach aggregating statistical and machine learning models generally achieved the best performance across all locations and forecast horizons. Relative forecast performance improved as the forecast horizon extended from 1 to 4 weeks ahead. The most challenging data to forecast were primarily restricted to low mosquito activity periods or atypical population peaks with unusual timing or magnitude. While full integration into routine vector management workflows represents a long-term process requiring operational adaptation, this work advances forecasting research and establishes a baseline for translating these approaches into real-time applications for public health authorities, with downstream effects in mitigating the risks of mosquito-borne diseases.

## Introduction

*Aedes aegypti* is a primary vector for several pathogens, such as dengue virus, chikungunya virus, and Zika virus, which cause millions of cases worldwide^1–5^. Within the continental United States, the public health landscape is defined by the dual pressure of established *Ae. aegypti* populations across the southern states^6,7^ and the frequent importation of these pathogens through international travel^8–10^. The convergence of these factors has triggered local transmission events, such as recent dengue outbreaks in Florida^11^, highlighting a growing risk that is expected to escalate over the coming years^12^.

The ability of local mosquito control authorities to mitigate *Aedes*-borne pathogen transmission during an outbreak depends on timely interventions such as larviciding and adulticiding^13,14^. By suppressing vector abundance ahead of an anticipated surge, vector control teams could lower human-mosquito contact rates, thereby reducing the probability that an imported case will initiate a localized transmission chain, thus preventing an outbreak from taking place. However, traditional vector management remains largely reactive, relying on contemporaneous surveillance data that reflects past or current population levels rather than future trends^15,16^. Transitioning to a proactive management strategy requires predictive tools that can anticipate surges in vector populations, thus allowing a better allocation of resources and personnel^17^.

This shift toward prospective information mirrors recent advancements in broader infectious disease management. The development of centralized forecasting hubs for influenza^18,19^, COVID-19^20^, and Respiratory Syncytial Virus^21^ by the Centers for Disease Control and Prevention (CDC), as well as similar international initiatives^22–24^, has shown the value of probabilistic forecasts in supporting public health decision-making^25^. However, this remains a partially unexplored area for mosquito populations^17^. Indeed, while most existing modeling work focuses on long-term trends and scenario analyses^12,26–29^, which provide essential ecological insights, these approaches lack the operational value of short-term forecasts.

The aim of this study is to develop and evaluate a multi-model ensemble framework designed to generate probabilistic 1-to 4-week-ahead forecasts of collected female *Ae. aegypti* per trap night. By integrating baseline time-series, statistical, and machine learning approaches, our ensembling approach aims to capture complex population dynamics while mitigating the risk of individual model limitations. Our analysis includes locations with highly divergent ecological and climatic profiles, utilizing time-series mosquito surveillance data from four US jurisdictions: Miami-Dade County, Florida; Key West, Florida; Los Angeles County, California; and Maricopa County, Arizona. Together, these study sites provide a comprehensive testbed for evaluating our forecasts across the varied environments (from subtropical to temperate and arid) where *Ae. aegypti* currently poses a public health threat.

## Methods

### Data

#### Mosquito Surveillance Data

We used mosquito surveillance data from four jurisdictions: For Key West, FL (2010-2020), Los Angeles County, CA (2018-2021), Maricopa County, AZ (2014-2021), and Miami-Dade County, FL (2017-2024). In all study sites, mosquitoes were collected by CO_2_-baited traps. Traps were deployed for 24 hours once a week in shaded areas protected from direct solar radiation, wind, and precipitation to enhance mosquito collection. Mosquitoes were morphologically identified to species using taxonomic keys. Trapping was conducted continuously throughout the calendar year for Key West, Maricopa County, and Miami-Dade County, while in Los Angeles County trapping efforts were based on a seasonal surveillance protocol. Traps baited with CO_2_ are intended to lure female mosquitoes in search of a blood meal. As a result, male mosquitoes collected during this project were deemed incidental catches and excluded from this study. All analyzed trap data were spatially aggregated at the jurisdictional level. This encompassed a cumulative pool of 12 traps in Key West, FL; 255 in Miami-Dade County, FL; 805 in Maricopa County, AZ; and 51 in Los Angeles County, CA, although the exact number of operational traps varied by week. To maintain a consistent temporal resolution across all study sites and mitigate the impact of irregular field schedules (e.g., a trap deployed on a Monday one week but a Tuesday the next), each epidemiological week was standardized to commence on Monday. Note that throughout this manuscript, “relative abundance per trap night” refers to the number of female *Ae. aegypti* collected per trap night in a given week.

#### Meteorological Data

To account for the environmental impact on *Ae. aegypti* development, mortality, and oviposition rates, as well as the carrying capacity of the environment^30,31^, we integrated average weekly temperature (°C) and total weekly precipitation (mm) into the modeling frameworks. Data for each study site were retrieved from Weather Underground (https://www.wunderground.com/).

### Forecasting framework

To forecast *Ae. aegypti* population dynamics, we developed a suite of statistical models. Forecasts were generated independently for each model for 1-to 4-week-ahead horizons. We also developed an ensemble approach that integrates the trajectories of the proposed models.

#### Training Method

To mimic real-world operational public health deployment, for any selected forecast date (i.e., the week at which a forecast is generated), the models were trained using the entire available dataset up to that week^25^. To ensure sufficient data for parameter estimation, we set a minimum of two full years of historical data; thus, no forecast dates were simulated within the first two years of any jurisdiction’s time series. For each subsequent week, the training dataset was expanded until the forecast week, and the models were recalibrated. Observed meteorological data (weekly temperature and precipitation) were used during the model training phases, while weather data during the forecast horizon were defined as the 5-year historical average calculated for each corresponding calendar week.

### Forecasting Method

#### Naïve Baseline

As a baseline, we implemented a “naïve” model whose forecasts are based on historical week-to-week differences, adopting a modified approach from CDC infectious disease forecasting initiatives^25^. For each week up to the forecast date, we calculated the difference in relative abundance from the preceding week. We then fit a normal distribution with a mean of zero and a standard deviation derived from these historical differences. Forecast trajectories were generated iteratively as *c*(*t*+1)=*c*(*t*)+ *ϵ*(*t*), where *ϵ*(*t*) is a value sampled from the fitted normal distribution and *c*(*t*) is the number of female *Ae. aegypti* collected per trap night for week *t*. For each study site and forecast date, we simulated 1,000 independent trajectories to construct the probabilistic, quantile-based forecasts.

#### Seasonal-Trend Decomposition using LOESS (STL)

We developed a weather-independent forecasting framework using additive Seasonal-Trend Decomposition based on LOESS (STL) coupled with stochastic Monte Carlo simulations to capture *Ae. aegypti* phenology. *Ae. aegypti* relative abundance per trap night in week *t* (where *t* belongs to the training period), *yt*, was decomposed as:

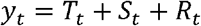

where:

- *c* represents the low-frequency trend capturing multi-year population dynamics at week *t*;
- *S*_*t*_ represents the 52-week seasonal component;
- *R*_*t*_ represents the remainder residual noise.

To account for the heteroscedasticity inherent in mosquito populations, residuals during the training period were stratified into two variance periods based on the sign of the seasonal component: a “high activity” period for weeks where *St* ≥0, and a “low activity” period for weeks where *S*_*t*_ <0. Future residual noise, *R*_*t*_, was sampled from a normal distribution *N*(0, *σ*^*2*^), using the specific standard deviation calculated for the corresponding activity period (i.e., *σ*_*high*_ or *σ*_*low*_).

For the 1-to 4-week forecast horizons, the baseline population level was set to the mean relative abundance per trap night over the preceding *n* weeks, where *n* ∈ {1,2,3,4}, yielding four distinct model variations (STL1 – STL4). From each of these four models, we generated 1,000 independent Monte Carlo trajectories. The relative abundance at each future step was estimated by applying the deterministic weekly change in the seasonal component combined with a randomly sampled residual. Then, 99 quantiles (ranging from 0.01 to 0.99) for each forecast horizon were extracted empirically from the distribution of the 1,000 simulated trajectories.

### Seasonal Autoregressive Integrated Moving Average with Exogeneous Factors (SARIMAX)

We used a Seasonal Autoregressive Integrated Moving Average extended with exogenous covariates (SARIMAX^32^) to capture the population dynamics and environmental dependencies of *Ae. aegypti*. The model was specified as a non-seasonal order of (1, 1, 2) and seasonal order of (1, 0, 0, 52). The first-order non-seasonal differencing was applied to stabilize the mean, while the autoregressive and moving average terms captured short-term population carryover and the smoothing of transient unobserved shocks. The seasonal autoregressive term explicitly modeled the 52-week annual cycle, and average weekly temperature and total precipitation were integrated as linear exogenous predictors.

Using the backshift operator *L* (where 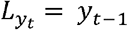), the SARIMAX model is defined as:

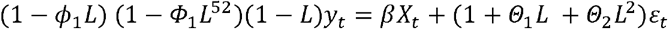

where:

- *y*_*t*_ is the relative abundance per trap night in week *t*;
- *ϕ*_1_ is the non-seasonal autoregressive coefficient that captures the carryover of the mosquito population from the previous week;
- *Ф*_*1*_ is the seasonal autoregressive coefficient accounting for annual phenology;
- (1−*L*)is the non-seasonal differencing operator;
- *β* is the linear regression weights for the exogenous predictors (i.e., average weekly temperature and total precipitation);
- *X*_*t*_ is the concurrent vector of average weekly temperature and total precipitation;
- *θ*_1_ and *θ*_2_ are the moving average coefficients modeling a two-week decay of transient shocks;
- *ε*_*t*_ ∼ *N*(0,*σ*^*2*^)is the unobserved ecological shocks embedded in the error term.

Seasonal differencing (*D*=0) was intentionally omitted to prevent over-differencing, as the model already incorporates non-seasonal differencing alongside seasonal exogenous drivers.

#### Unobserved Components Model (UCM)

We implemented an Unobserved Components Model (UCM) that explicitly formulates the observed time series as the linear combination of independent evolving components:

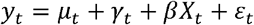

where:

- *y*_*t*_ is the relative abundance per trap night in week *t*;
- *μ*_*t*_ is a random walk that adapts to shifts in *Ae. aegypti* relative abundance;
- *γ*_*t*_ is the 52-week seasonal component that captures annual cyclicity;
- *βX*_*t*_ is the linear regression effect of the exogenous predictors (i.e., average weekly temperature and total precipitation);
- *ε*_*t*_ is a Gaussian white noise error.

#### Ensemble Model

Our ensemble model was constructed by combining the probabilistic outputs of six constituent models: SARIMAX, UCM, and the four STL variations. The naïve baseline was excluded from this aggregation. For each jurisdiction, forecast date, and forecast horizon, the ensemble’s predicted quantiles were calculated as the mean of the corresponding quantiles from the constituent models.

For a given quantile *q* at forecast horizon *h*, the ensemble quantile *Q*_*ens*_ (*q*)was defined as:

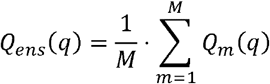

where *M* is the number of constituent models (i.e., *M* = 6) and *Qm* (*q*) represents the quantile projection from model *m*.

This averaging was performed simultaneously across all 99 discrete quantiles to generate the final probabilistic distribution.

### Forecast Evaluation

#### Probabilistic Quantile Generation

For each location and forecast horizon, we generated 99 discrete quantiles ranging from 0.01 to

0.99. For the parametric models (SARIMAX and UCM), these quantiles were estimated by assuming a Gaussian distribution of the forecast errors:

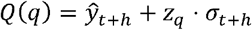

Where *ŷ*_*t+h*_ is the predicted mean at horizon *h, σ*_*t*+*h*_ is the standard error of the forecast at horizon *h*, and *z*_*q*_ is the standard normal inverse cumulative distribution function for quantile *q*.

To maintain biological validity, any resulting quantile falling below zero was bounded at zero.

#### Forecast Evaluation Metrics

To assess model performance, we used the Weighted Interval Score (WIS), a weighted average of the performance of each prediction interval that assesses both the precision and accuracy of a forecast, which is used as the main metric to evaluate forecast performance during CDC infectious disease forecast initiatives^25^. The WIS evaluates the entire predictive distribution by penalizing models for generating excessively wide uncertainty intervals (lack of sharpness) and for poor calibration (when the actual observed mosquito counts repeatedly fall outside the predicted confidence intervals). A lower WIS indicates better performance.

For any given target week *t*, the absolute WIS is calculated across *K* central prediction intervals(e.g., 0.1, 0.5, 0.9) and the model’s forecasted median (*m*_*t*_) as:

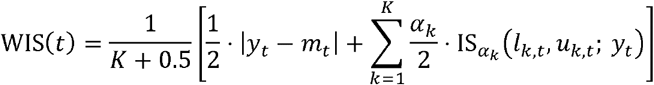

where:

- *y*_*t*_ is the *Ae. aegypti* relative abundance per trap night at week *t*;
- *m*_*t*_ is the model’s forecasted median at week *t*;
- *α*_*k*_ is the nominal probability that an observation falls outside the *k*-th prediction interval(e.g., *α*_*k*_ =0.1 for a 90% prediction interval).
- 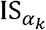 is the interval score for the *k*-th prediction interval, which is calculated as:

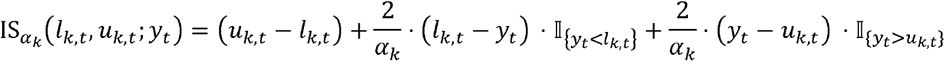

where *l*_*k*_,_*t*_, and *u*_*k*_,_*t*_ are the lower and upper bounds of the prediction interval *k* at week *t*, respectively; I_{expression}_ is an indicator function equal to 1 if the expression is true and 0otherwise. Note that the first term, *u*_*k*_,_*t*_ − *l*_*k*_,_*t*_, measures the width of the interval, rewarding narrower forecasts, while the second and third terms apply a penalty of 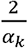 when the observed value falls outside the forecasted interval.

To directly compare model performance against the naïve baseline, we calculated the Relative WIS (*rWIS*) for each model as the ratio of its absolute WIS to the WIS of the naïve baseline across the same target weeks:

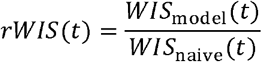

An *rWIS* < 1indicates that the model outperformed the naïve baseline.

To evaluate forecast performance, we added two additional statistical metrics:

- sMAPE: The model’s forecasted median at week was used to calculate the Symmetric Mean Absolute Percentage Error (sMAPE)^17^ to evaluate point accuracy:

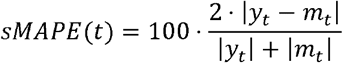

where the notation is the same used in the definition of WIS. This metric is symmetric between the forecast and observation, it is bound between 0 and 200, and lower values representing more accurate forecasts.
- Coverage: This metric measures the proportion of the observed values that fall within the forecasted upper and lower bounds^25^, and it is calculated as:

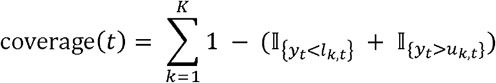

where the notation is the same used in the definition of WIS.

## Results

Figure 1 shows a visual assessment of individual models’ forecasts against one year of mosquito surveillance data. Miami-Dade County, FL, and Maricopa County, AZ, were selected as illustrative examples to compare forecast performance across different ecological profiles (subtropical and arid, respectively). Forecasts for every analyzed year and location are reported in the Supplementary Material (Fig. S1-S28). A visual inspection of the individual constituent models shows two distinct behaviors over the year (Fig. 1A): during low mosquito activity periods (e.g., late fall to early spring), all models yield comparable quantitative forecasts; conversely, during peak mosquito activity periods, the models exhibit noticeable variations in their forecasted medians, reflecting their distinct underlying model structures. Throughout the year, none of the individual models generate biologically implausible forecast intervals, with upper bounds in line with observed peak relative abundance per trap night. Aggregating individual model forecasts into the multi-model ensemble smooths out individual models, with observed data falling within the 50% prediction interval of the ensemble forecast for most of the year in both locations (Fig. 1B).

**Figure 1.**
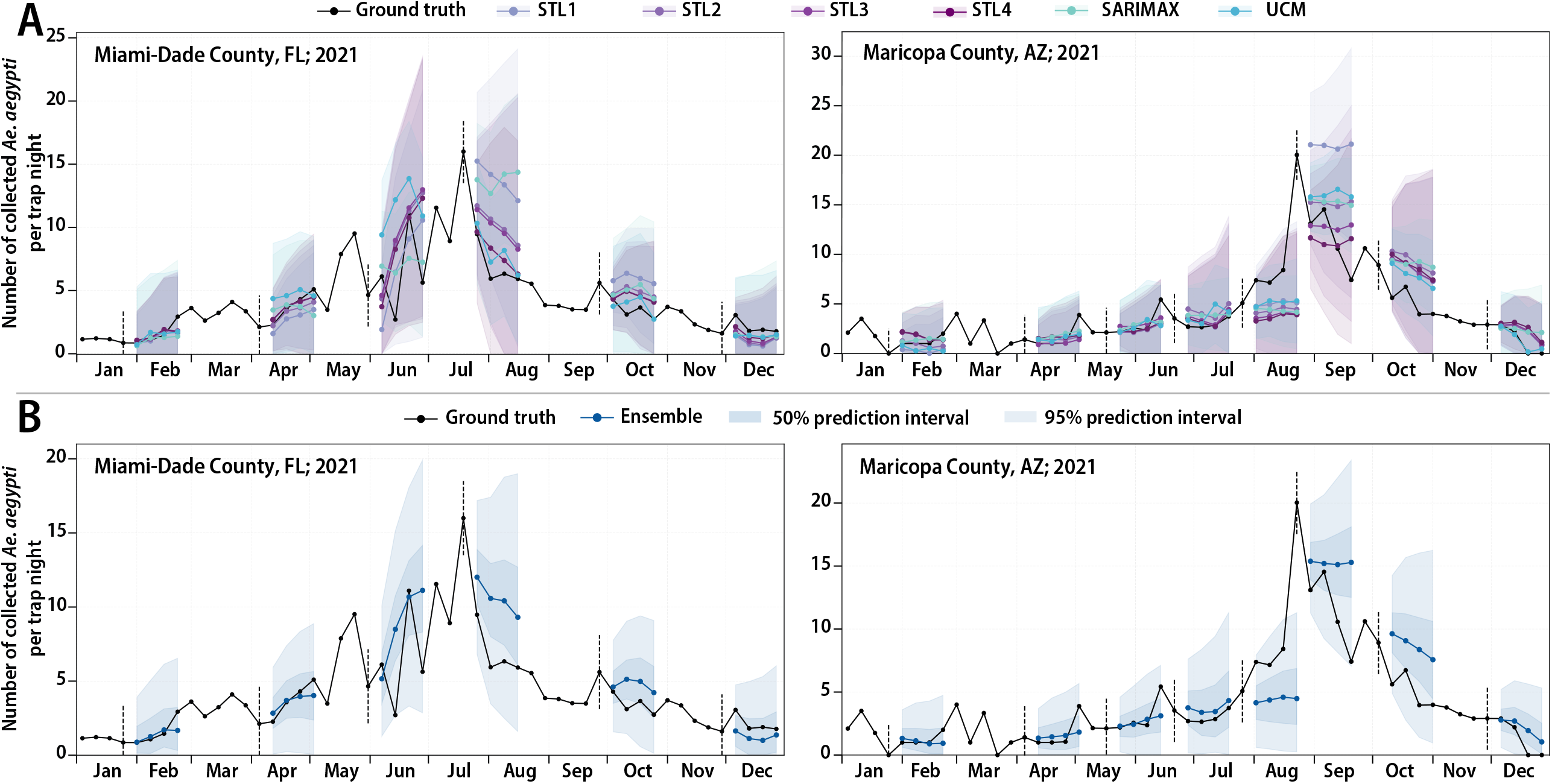
**A.** Comparison between the observed *Ae. aegypti* relative abundance per trap night in Miami-Dade County, FL, and Maricopa County, AZ, and individual model 4-week-ahead forecasts (median and 95% prediction interval) for a set of forecast dates. **B**. Comparison between the observed *Ae. aegypti* relative abundance per trap night in Miami-Dade County, FL, and Maricopa County, AZ, and 4-week-ahead forecasts from the ensemble model (median, 50% and 95% prediction interval) for a set of forecast dates.

Figure 2 departs from visual inspection and presents statistical metrics for all study sites and across all the available years of data. The relative performance of the models, evaluated via the Relative Weighted Interval Score (rWIS) against the naïve baseline, shows distinct patterns across study locations. Across all forecast horizons, the lowest rWIS was recorded for Los Angeles County, followed by Miami-Dade County and Key West, while the weakest performance was observed in Maricopa County (Fig. 2).

**Figure 2.**
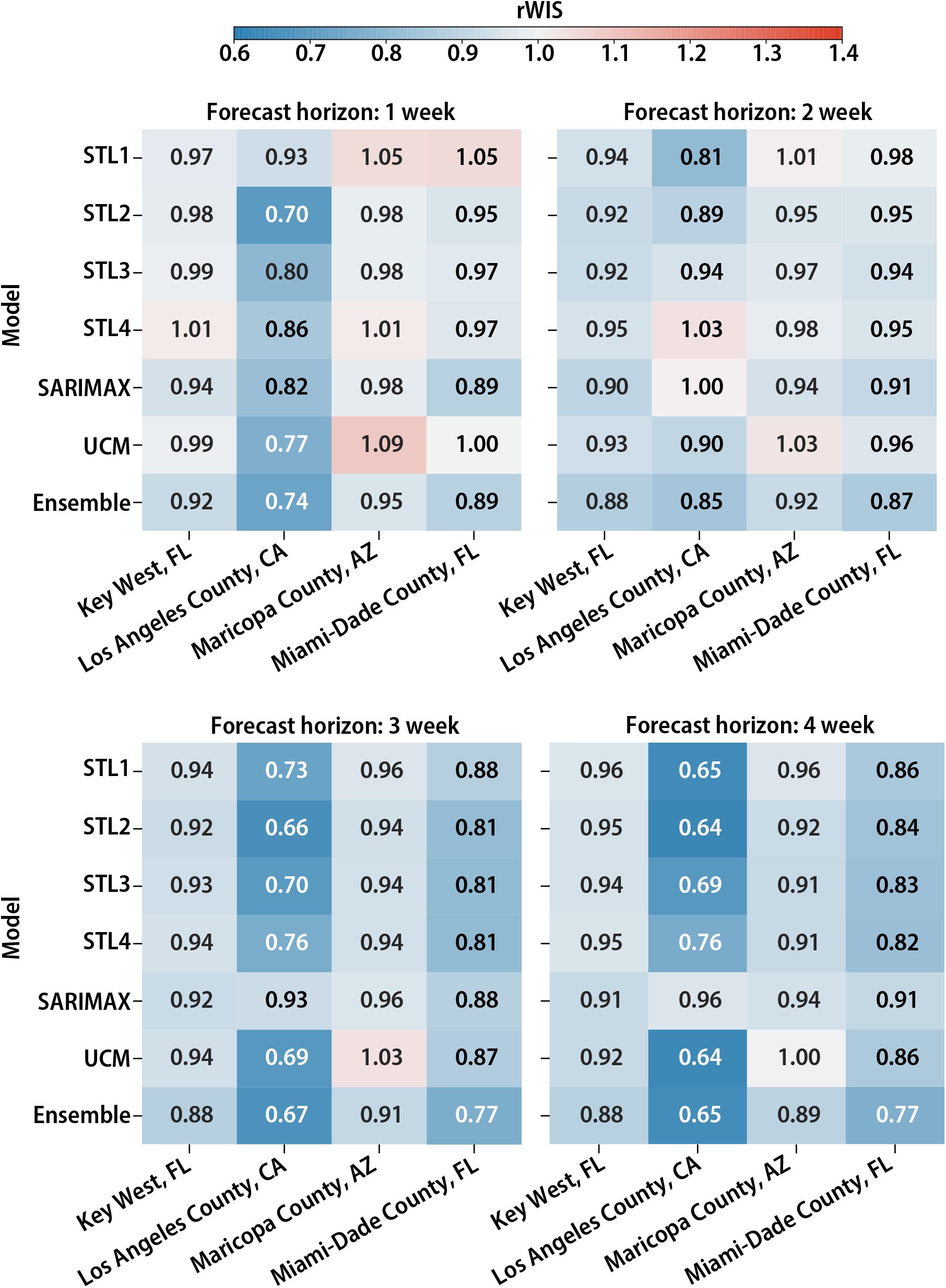
Heatmaps reporting the relative WIS estimated for each independent model and the ensembled forecast for each location and forecast horizon. This figure is generated using all years of available mosquito surveillance data.

Forecast performance varied by model (Fig. 2). The seasonal decomposition models (STL1 to STL4) show a performance gradient, where shorter historical baseline windows (*n* = 1 or 2) generally yield stronger forecasts than longer averages. The SARIMAX model appears to be conservative, consistently slightly overperforming the naïve model across most locations and time horizons (rWIS between 0.89 and 1 in 15 out of 16 scenarios). The UCM model shows a higher performance variability (rWIS between 0.64 and 1.09). Aggregating individual model forecasts into the multi-model ensemble smooths out individual model variations and provides consistently good performance (rWIS < 0.9 in 12 out of 16 scenarios). Moreover, the ensemble model of leads to better forecasts than any single model, including in the hardest scenario considered: the 1-week-ahed forecasts in Maricopa County where the ensemble model yielded a rWIS of 0.95, while no single model scored a value lower than 0.98. Similar patterns were found when considering other forecasts evaluation metrics (see Table S1-S4 in the Supplementary Material).

Unlike traditional public health forecasting efforts^22,25^ where forecast performance decreases for longer time horizons, our forecast performance is actually improving (Fig. 2). Across all locations, the ensemble model shows a consistent decrease in rWIS from 1-to 4-week-ahead forecasts, with 4-week-ahead forecasts performing best, followed by 3-, 2-, and 1-week-ahead forecasts.

Forecast performance was highly variable between epidemiological weeks (Fig. 3). Periods of the year when models underperformed relative to the baseline (i.e., rWIS > 1) typically corresponded to two categories. The first category corresponds to periods of the year where one or more historical years experienced a mosquito population peak with unusual magnitude or atypical timing (as it was the case in Miami-Dade County, Maricopa County, and Key West). The second category corresponds to periods of very low mosquito activity, where the number of collected mosquitoes per trap night is near-zero (e.g., December to April in Key West and the very start of the season in Los Angeles County).

**Figure 3.**
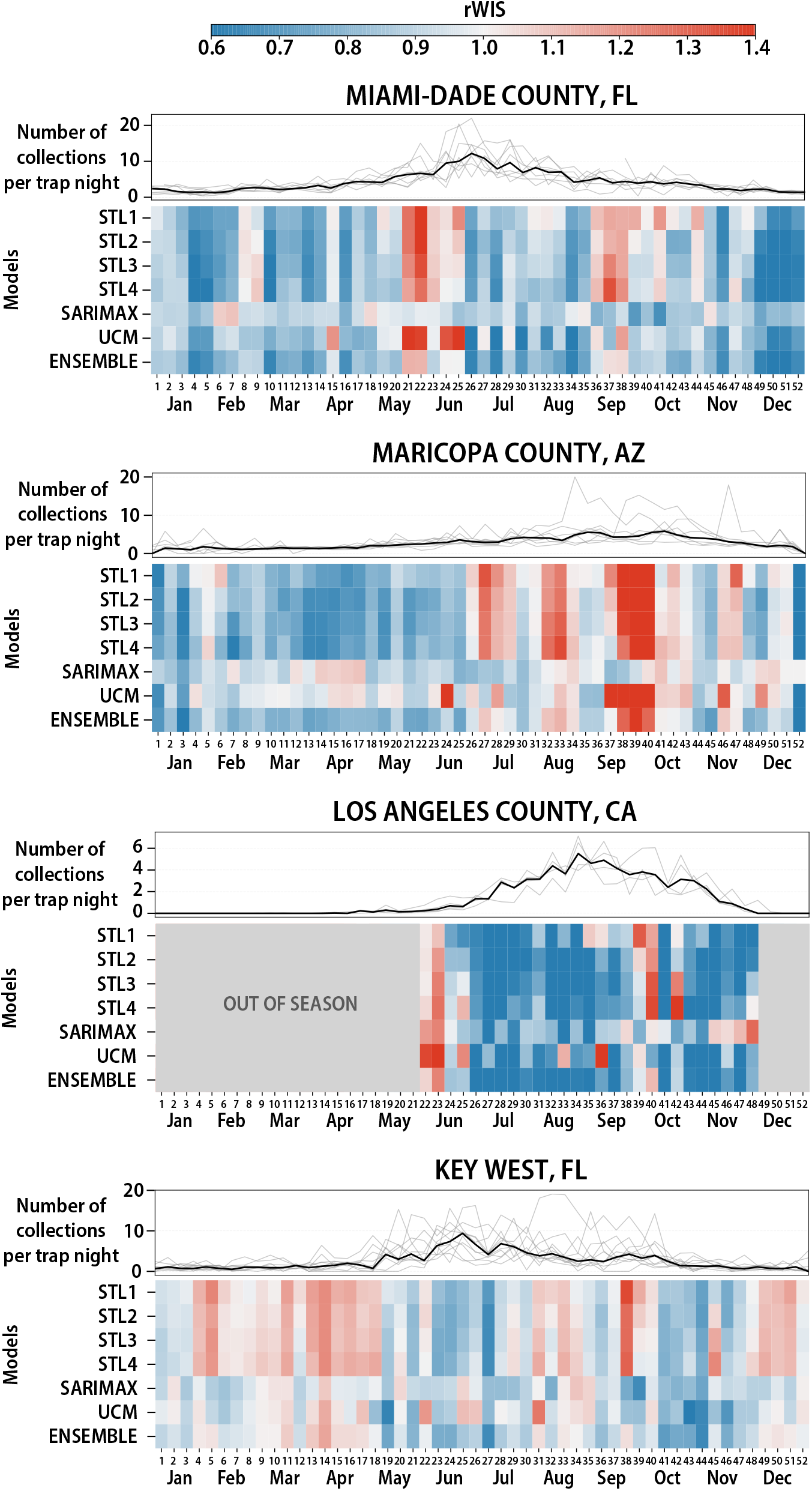
Heatmaps of the relative WIS estimated for each independent model and the ensembled forecast for each location and epidemiological week. The estimated rWIS for each week represents the average on that week across all the available years of date for each location. On top of each heatmap, the line plot shows *Ae. aegypti* relative abundance per trap night for each year of data (grey lines). The black line represents the median across years.

This longitudinal breakdown also highlights substantial variability between the individual models (Fig. 3). The STL models provided generally good forecasts relative (rWIS < 1) with localized periods of lower performance. SARIMAX’s forecasts were highly conservative, consistently hoovered close to the naïve baseline model across almost all weeks. UCM showed more variability than SARIMX, but lower performance compared to the STL models. The ensemble model provided the most reliable forecasts throughout the year, outperforming the baseline model almost every week for all locations.

## Discussion

This study evaluated a suite of statistical approaches to generate 1-to 4-week-ahead probabilistic forecasts of female *Aedes aegypti* relative abundance per trap night. By assessing these models across four US jurisdictions, the framework was validated under highly different environmental conditions, from arid to subtropical and temperate. Our results show that using an ensemble approach generally yield the best performing forecasts across any location and forecast horizon. Forecast performance relative to the baseline model improved as the forecast horizon extended from 1 to 4 weeks ahead. This pattern contrasts with many infectious disease forecasting applications where accuracy degrades over longer horizons^22,25^, suggesting that the proposed forecasting framework could be useful for operational vector control planning.

Although our ensemble approach achieved the best performance, it relied on a simple unweighted quantile averaging method. This framework assigns an equal weight to each constituent model, a baseline strategy frequently adopted in collaborative public health forecasting initiatives^25^. In this study, this approach is based on the aggregation of six models, four of which are variations of the same seasonal-trend decomposition framework. While distinct in their parameterizations, these models are conceptually similar and collectively account for two-thirds of the ensemble formulation. This structural configuration suggests that there is substantial potential to improve forecast performance by implementing more advanced ensembling techniques and/or increasing the number of ensembled models. For instance, instead of simple averaging, weighted combinations, performance-based weighting, meta-learning, or voting techniques can be utilized to dynamically adjust model contributions^33,34^. Assessing these alternative ensembling methods represents a promising future direction to improve our forecasting framework.

When evaluated individually, the proposed forecasting models showed distinct characteristics across the time series. SARIMAX generated conservative forecasts that closely tracked the naïve baseline. Conversely, the seasonal-trend decomposition models performed well in jurisdictions with regular annual cycles but had lower performance in locations with high interannual variability in peak abundance (e.g., Maricopa County). Beyond regional seasonality, the performance of these models across forecast horizons was sensitive to the scale of data aggregation and local stochastic noise. In jurisdictions with a lower number of traps, such as Key West (12 traps vs. 51, 255, and 805 traps in Los Angeles, Miami-Dade, and Maricopa counties, respectively), the mosquito surveillance data is subject to higher week-to-week volatility. Because the seasonal-trend decomposition frameworks structurally prioritize low-frequency historical trends, they filter out this high-frequency transient noise, contributing to more stable performance over longer horizons compared to larger jurisdictions where the heavily aggregated signal is already smoothed. Moreover, since the initialization of STL models is sensitive to the relative abundance observed immediately prior to the forecast date, integrating nowcasting tools to establish the baseline population level could represent an important research direction to improve model performance.

From an operational perspective, the most challenging phases for our forecasting framework fell into two categories: i) low mosquito activity periods, and ii) periods characterized by atypical population peaks, where surges occurred with unusual timing or magnitude. Low mosquito activity periods have limited interest from an operational perspective, as mosquito control interventions are generally minimal when vector abundance is near zero. Conversely, a reduction in forecast performance during periods characterized by atypical population peaks carries high operational relevance because these are windows where proactive intervention could be critical. This operational challenge varied by location, and it was visible in Maricopa County, while it appeared to be less relevant in the other three locations. Our forecasting framework’s difficulty in consistently capturing these peaks underscores the need for model improvements. Another key consideration for the operational utility of the forecasts is aligning their resolution with the spatial scales where mosquito control authorities typically target interventions, such as specific neighborhoods. Although this study was conducted at the jurisdictional level, multiple individual neighborhoods within Miami-Dade County and Maricopa County already maintain 10 or more active traps, which aligns with the total number of traps deployed across Key West. Because the forecasting framework achieved stable performance in Key West under this data constraint, there is a rationale that our framework could operate effectively when deployed at the neighborhood level. Therefore, evaluating our framework at finer spatial resolution represents a critical direction for future development to enhance the operational utility of the forecasts.

This study is subject to several limitations. First, because the analysis was conducted at the jurisdictional level, environmental covariates for SARIMAX and UCM were restricted to temperature and precipitation, which show clear temporal fluctuations at this scale. Other drivers that influence mosquito population dynamics through spatial variation, such as vegetation and land use, could not be integrated due to the spatial aggregation of the surveillance data used for this study. Second, the parameters and lag structures for the SARIMAX model were selected based on established relationships reported in the literature rather than through formal statistical optimization tests to identify best-fit configurations. Third, during the 1-to 4-week forecast horizons, our framework relies on a 5-year historical average to represent temperature and precipitation rather than integrating real-time meteorological forecasts. While this approach provides a stable baseline for operational planning, it inherently limits the models’ capacity to anticipate sudden, unseasonal weather anomalies such as severe heatwaves or heavy rainfall events, which could alter short-term *Ae. aegypti* population dynamics and temporarily hamper real-time forecast performance. Fourth, our study focused on a single vector species across four specific geographic areas. These findings may thus not directly generalize to other mosquito species, distinct ecological regions, or alternative spatial scales. Fifth, our forecasting framework generates probabilistic forecasts rather than categorical classifications, such as defining trends as increasing, stable, or decreasing. Consequently, forecast evaluation was restricted to quantitative metrics (primarily the WIS and additional metrics such as sMAPE and coverage). While probabilistic distributions preserve full uncertainty information, operational public health decisions often rely on categorical action thresholds. Adopting a categorical forecasting approach would require establishing and validating these specific thresholds, which introduces distinct verification challenges that were outside the scope of this study.

While the primary focus of this work is to advance research in mosquito population forecasting, it also establishes a baseline for translating time-series models into real-time applications for public health authorities. Transitioning this forecasting framework into a fully integrated management system represents a long-term process that requires future investments. Ultimately, once successfully implemented within routine, the decision-making workflows of mosquito control authorities, this proactive approach could contribute to limiting vector abundance, leading to downstream epidemiological effects in lowering the risk of mosquito-borne diseases.

## Code Availability

Models’ parameters are reported in the Supplementary Material and full codebase to reproduce all the experiments is openly available on GitHub at https://github.com/CEPH-Lab/ae_aegypti_forecasting.

## Data Availability

Data from Key West, Florida, is publicly available at Pruszynski et al.^36^. Data from Miami-Dade, Maricopa, and Los Angeles counties analyzed in this study were obtained from publicly supported surveillance programs. Although such data are public property and may ultimately become part of the public record, access is not automatically unrestricted or immediate. Individuals seeking access to the surveillance data used in this study must obtain permission from the appropriate public health authority to ensure compliance with applicable public-records statutes, data-use policies, and privacy protections. Researchers interested in obtaining the dataset may submit a formal Public Records Request to:

- Miami-Dade County Mosquito Control Division, available at: https://www.miamidade.gov/global/solidwaste/mosquito/contact-mosquito-control.page
- Maricopa County Environmental Services, Department of Vector Control Division, available at: https://publicrecordsrequest.maricopa.gov/nonCommercial?d=133
- Greater Los Angeles County Vector Control District, through the California surveillance data request form available at https://vectorsurv.org

## Acknowledgements

The authors would like to thank Susanne Kluh and Steve Vetrone from the Greater Los Angeles County Vector Control District, Santa Fe Springs, CA, USA, for providing access to mosquito surveillance data for Los Angeles County and supporting this research project.

