## Supplementary Material for "Short-term forecasts of *Aedes aegypti* relative abundance to enhance mosquito control situational awareness"

Utkarsh Bhosekar[1], Paulo C. Ventura[1], Megan D Hill[1], Allisandra G. Kummer[1], Shreeya Mhade[1], Jagadeesh Chitturi[1], Chalmers Vasquez[2], John-Paul Mutebi[2], John Townsend[3], Maria Litvinova[4], Andre B.B. Wilke[4], Marco Ajelli[1,*]

[1] Laboratory of Computational Epidemiology and Public Health, Department of Epidemiology and Biostatistics, Indiana University School of Public Health, Bloomington, IN, USA

[2] Miami-Dade County Mosquito Control Division, Miami, Florida, USA

[3] Maricopa County Environmental Services, Department Vector Control Division, Phoenix, AZ, USA

[4] Department of Epidemiology and Biostatistics, Indiana University School of Public Health, Bloomington, IN, USA

### S1. Model parameters

#### *S1.1. SARIMAX model*

The SARIMAX model uses sets of seasonal and trend parameters. The seasonal parameters are defined as $(P, D, Q, s) = (1,0,0,52)$, where $P$ is seasonal autoregressive terms, $D$ is seasonal differencing order, $Q$ is moving average terms, and $S$ is seasonal period (weeks). The trend parameters are defined as $(p, d, q) = (1,1,2)$, where $p$ is number of autoregressive terms, $d$ is order of differencing, and $q$ is number of moving average terms. Steps in the model were set to a value of 4 to estimate 1- to 4-week ahead forecasts. Exogenous predictors used for training the model included mean weekly temperature and total precipitation. Exogenous predictors used for the forecasting were a rolling 5-year average of the weekly temperature and total precipitation per ISO week. We set the maximum number of iterations for the optimizer to 50.

#### *S1.2. UCM model*

The UCM model uses a trend component that was set to “local level”, which acts as a random walk, and a seasonal component with a 52-week period for annual periodicity. As was used in the SARIMAX model, the exogenous predictors used for training the model included mean weekly temperature and total precipitation, an exogenous predictors used for the forecasting were a rolling 5-year average of the weekly temperature and total precipitation per ISO week.

#### *S1.3. Seasonal-trend decomposition models*

For the seasonal-trend decomposition models, we use one-sided, additive models as this method only uses past data until the training date for the forecast and separates the trend, seasonality, and residual noise. The seasonal period of the model was set to 52 weeks with the number of past weeks used to calculate the starting point was set to either 1, 2, 3, or 4, corresponding to each model used. We used two standard deviations for the residual noise added to the forecast which was categorized based on whether the forecast was being conducted for a high or low activity period ($\sigma_{high}$ and $\sigma_{low}$). These models were run for 1,000 iterations.

### S2. Additional results

#### *S2.1. Observed vs. Forecasted Aedes aegypti relative abundance*

We produce forecasts for weekly horizons between 1 and 4 weeks ahead of each reference date. Here we show plots for 1-, 2-, 3- and 4-weeks horizons median and 90% IQR forecasts over time. Forecasts for each individual model and the ensemble model in each location and for the 1- to 4-week ahead forecasts are shown in Figures S1-S28.

##### S2.1.1 STL1 model forecast

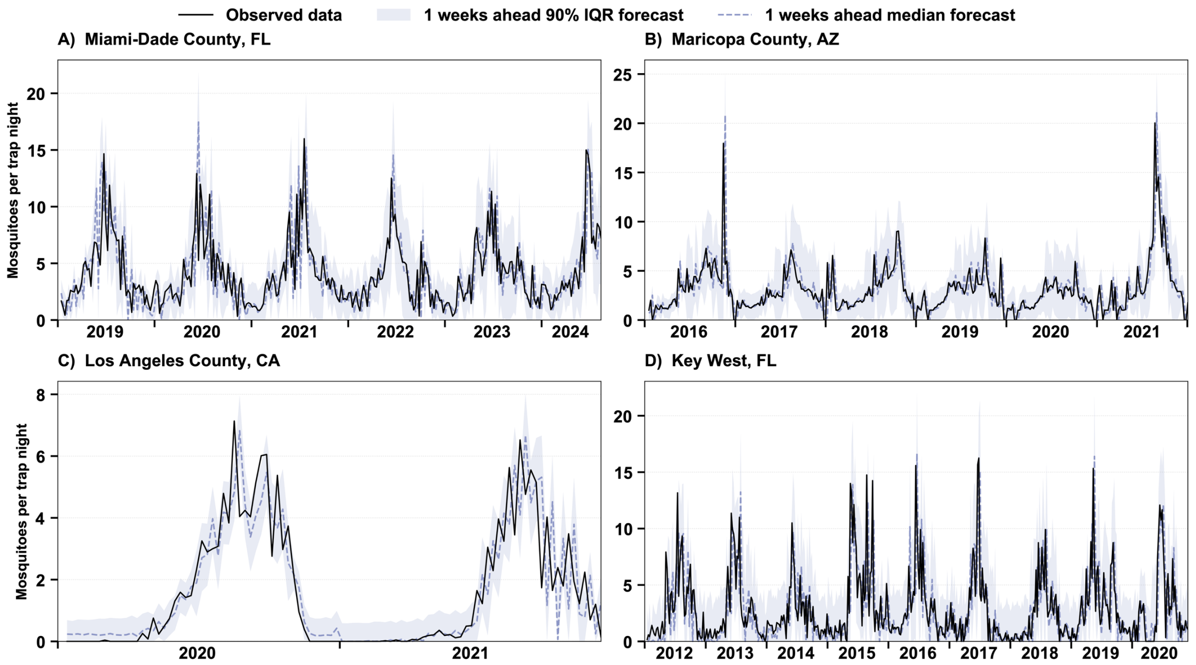

**Figure S1. Comparison between the observed *Ae. aegypti* relative abundance per trap night and 1-week-ahead forecasts (median and 90% prediction interval) for all forecast dates for the STL1 model. A.** Miami-Dade County, FL. **B.** Maricopa County, AZ. **C.** Los Angeles County, CA. **D.** Key West, FL.

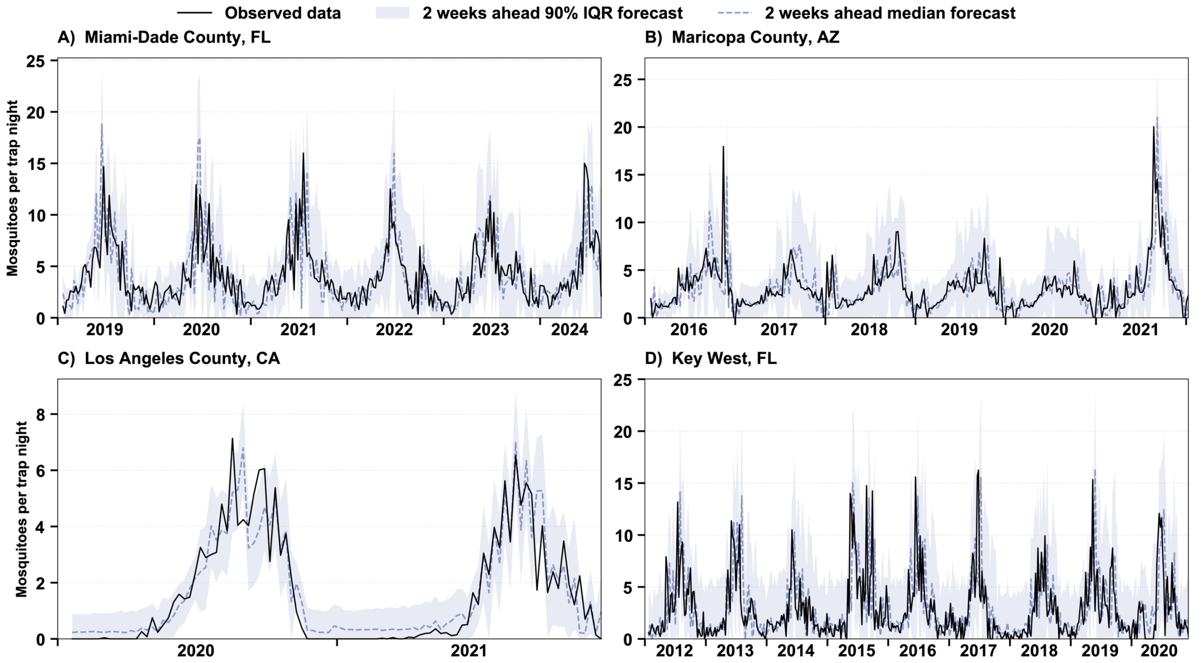

**Figure S2. Comparison between the observed *Ae. aegypti* relative abundance per trap night and 2-week-ahead forecasts (median and 90% prediction interval) for all forecast dates for the STL1 model. A.** Miami-Dade County, FL. **B.** Maricopa County, AZ. **C.** Los Angeles County, CA. **D.** Key West, FL.

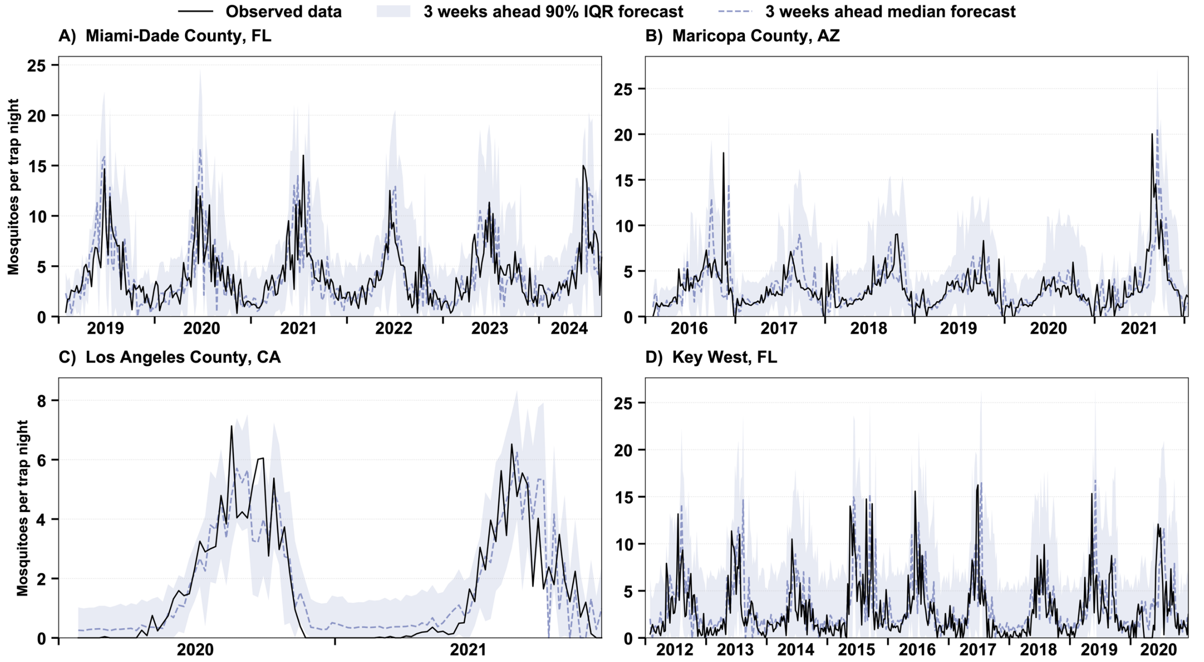

**Figure S3. Comparison between the observed *Ae. aegypti* relative abundance per trap night and 3-week-ahead forecasts (median and 90% prediction interval) for all forecast dates for the STL1 model. A.** Miami-Dade County, FL. **B.** Maricopa County, AZ. **C.** Los Angeles County, CA. **D.** Key West, FL.

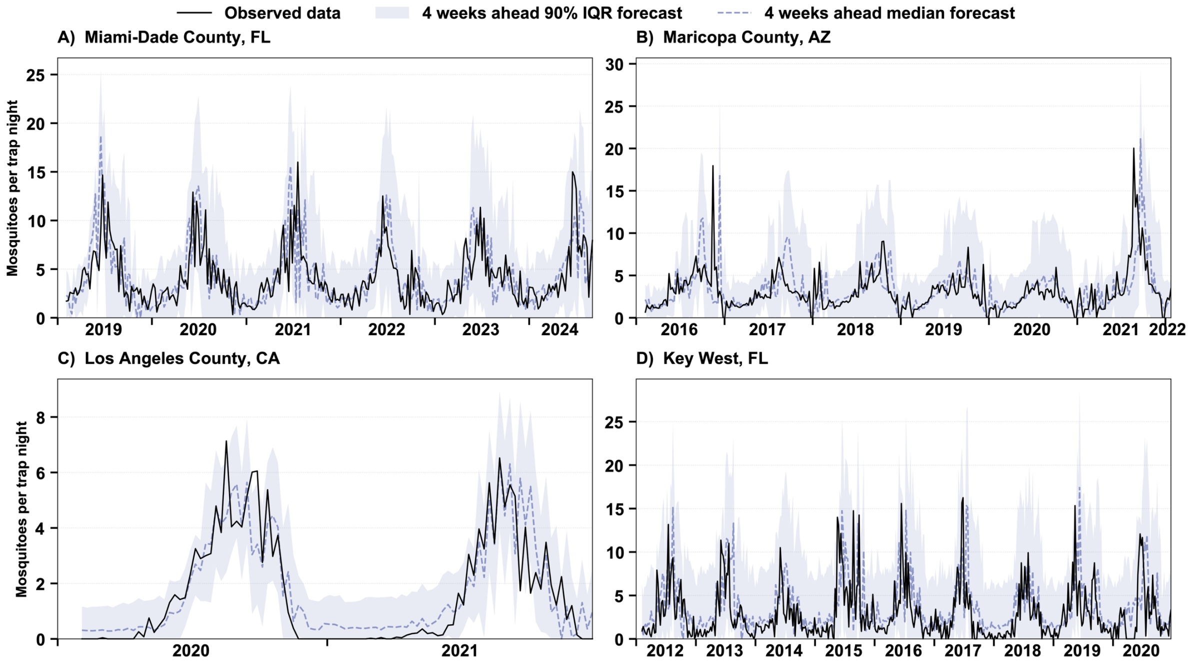

**Figure S4. Comparison between the observed *Ae. aegypti* relative abundance per trap night and 4-week-ahead forecasts (median and 90% prediction interval) for all forecast dates for the STL1 model. A.** Miami-Dade County, FL. **B.** Maricopa County, AZ. **C.** Los Angeles County, CA. **D.** Key West, FL.

### S2.1.2 STL2 model forecast

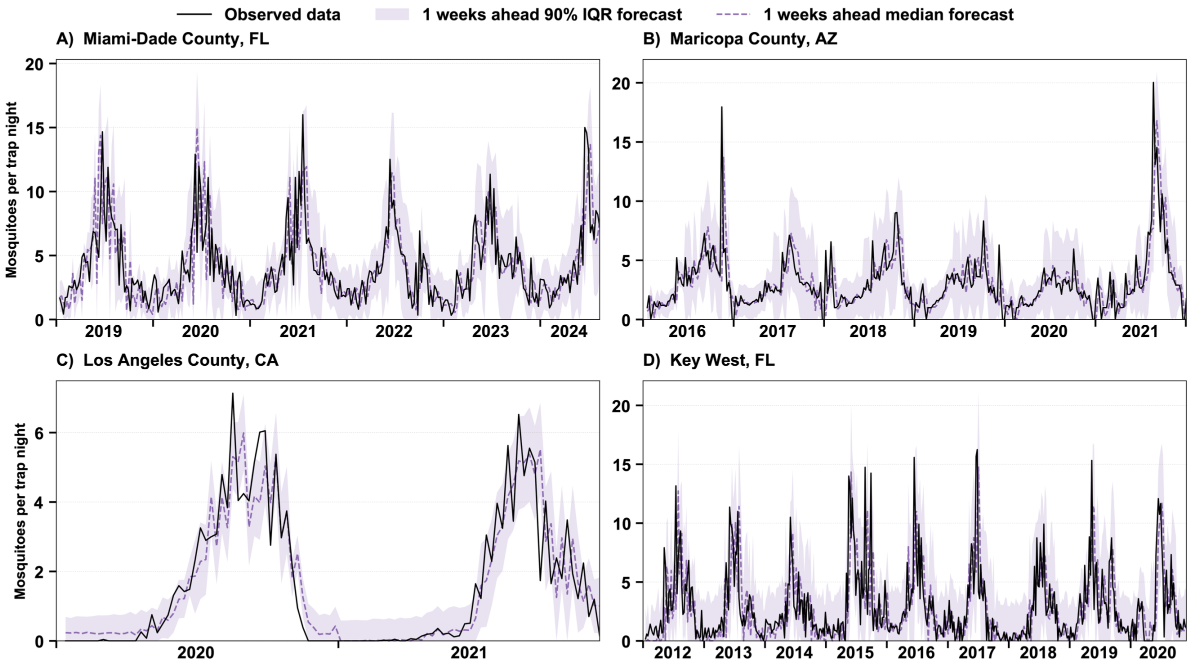

**Figure S5. Comparison between the observed *Ae. aegypti* relative abundance per trap night and 1-week-ahead forecasts (median and 90% prediction interval) for all forecast dates for the STL2 model. A.** Miami-Dade County, FL. **B.** Maricopa County, AZ. **C.** Los Angeles County, CA. **D.** Key West, FL.

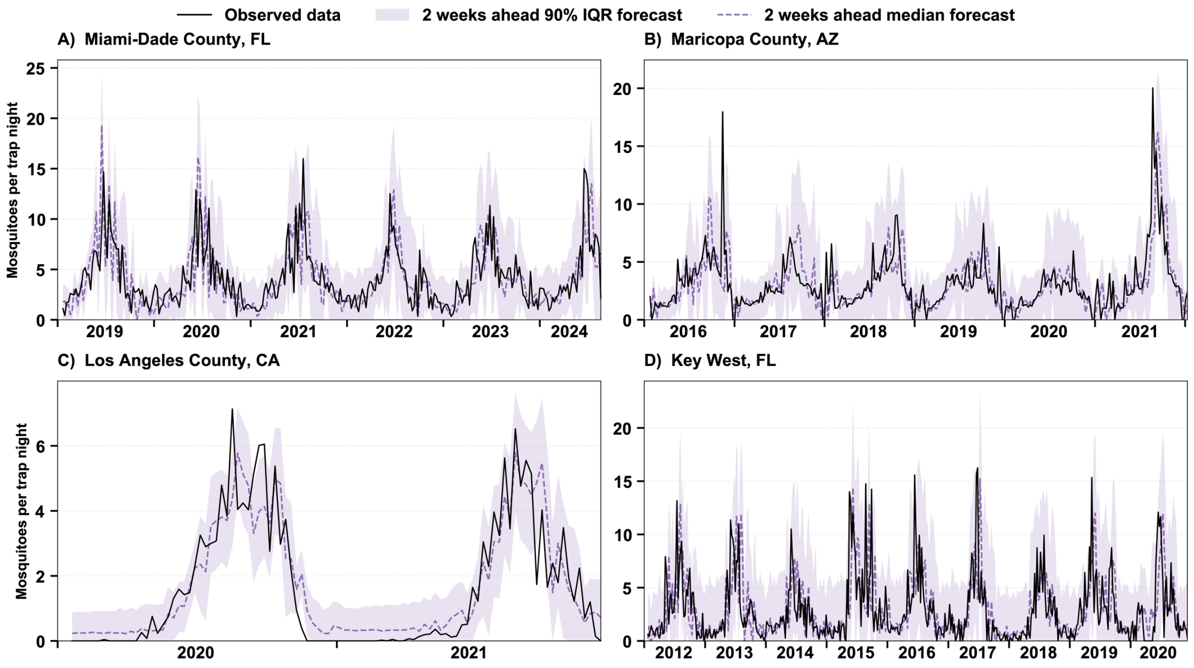

**Figure S6. Comparison between the observed *Ae. aegypti* relative abundance per trap night and 2-week-ahead forecasts (median and 90% prediction interval) for all forecast dates for the STL2 model. A.** Miami-Dade County, FL. **B.** Maricopa County, AZ. **C.** Los Angeles County, CA. **D.** Key West, FL.

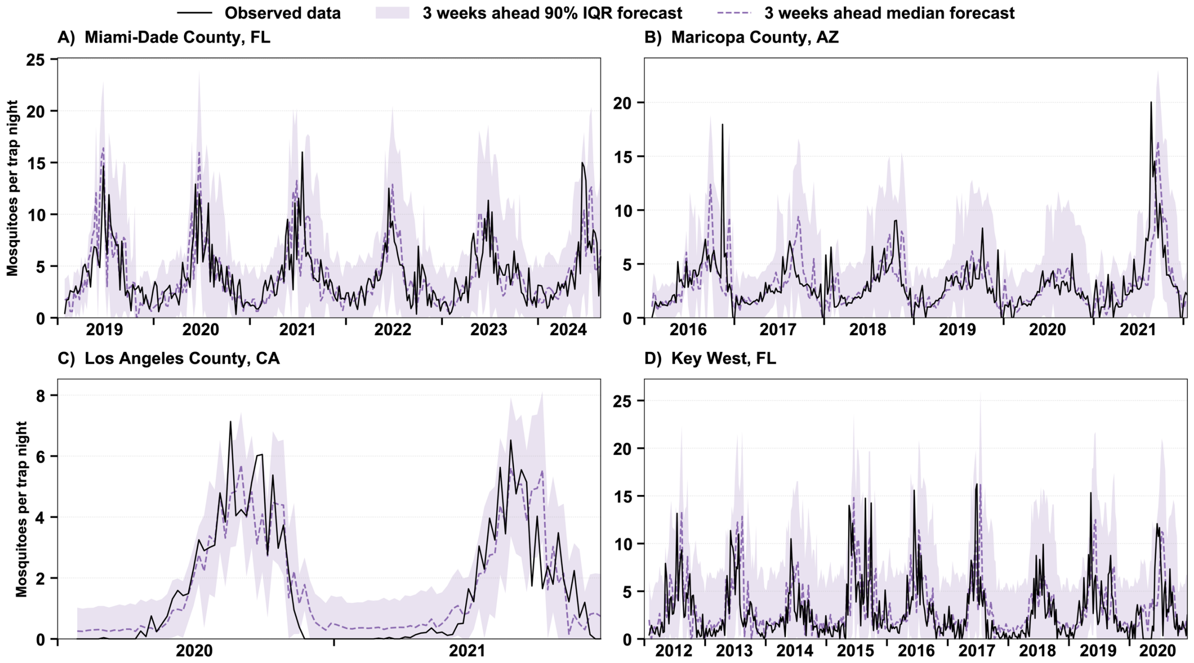

**Figure S7. Comparison between the observed *Ae. aegypti* relative abundance per trap night and 3-week-ahead forecasts (median and 90% prediction interval) for all forecast dates for the STL2 model. A.** Miami-Dade County, FL. **B.** Maricopa County, AZ. **C.** Los Angeles County, CA. **D.** Key West, FL.

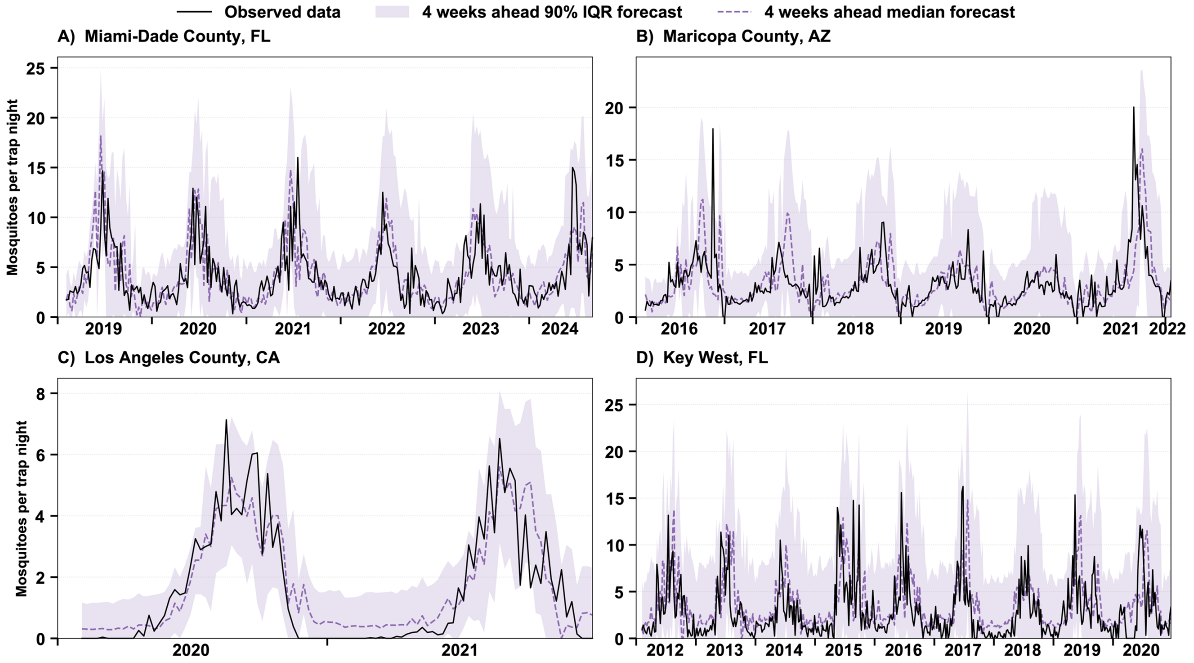

**Figure S8. Comparison between the observed *Ae. aegypti* relative abundance per trap night and 4-week-ahead forecasts (median and 90% prediction interval) for all forecast dates for the STL2 model. A.** Miami-Dade County, FL. **B.** Maricopa County, AZ. **C.** Los Angeles County, CA. **D.** Key West, FL.

### S2.1.3 STL3 model forecast

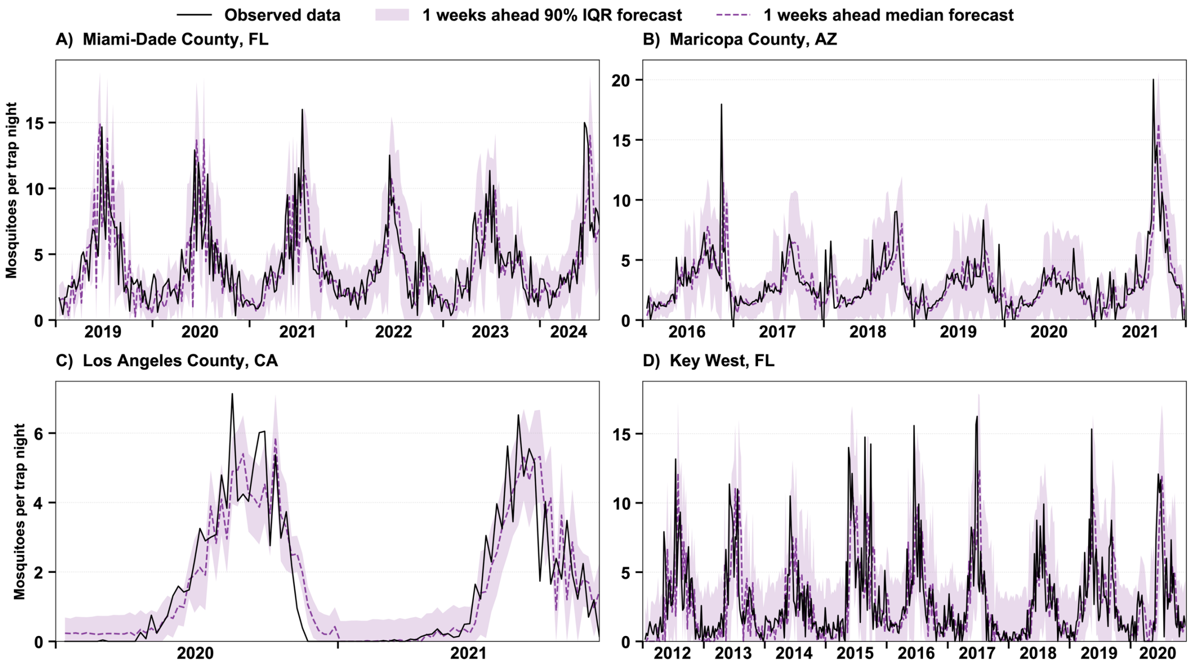

**Figure S9. Comparison between the observed *Ae. aegypti* relative abundance per trap night and 1-week-ahead forecasts (median and 90% prediction interval) for all forecast dates for the STL3 model. A.** Miami-Dade County, FL. **B.** Maricopa County, AZ. **C.** Los Angeles County, CA. **D.** Key West, FL.

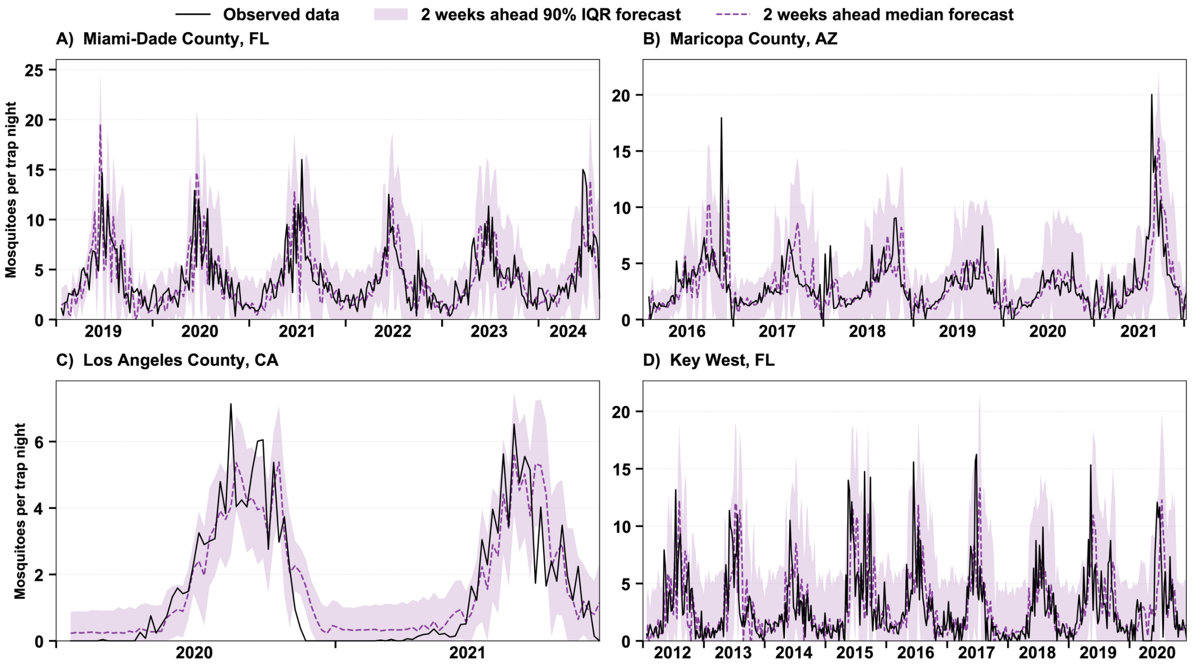

**Figure S10. Comparison between the observed *Ae. aegypti* relative abundance per trap night and 2-week-ahead forecasts (median and 90% prediction interval) for all forecast dates for the STL3 model. A.** Miami-Dade County, FL. **B.** Maricopa County, AZ. **C.** Los Angeles County, CA. **D.** Key West, FL.

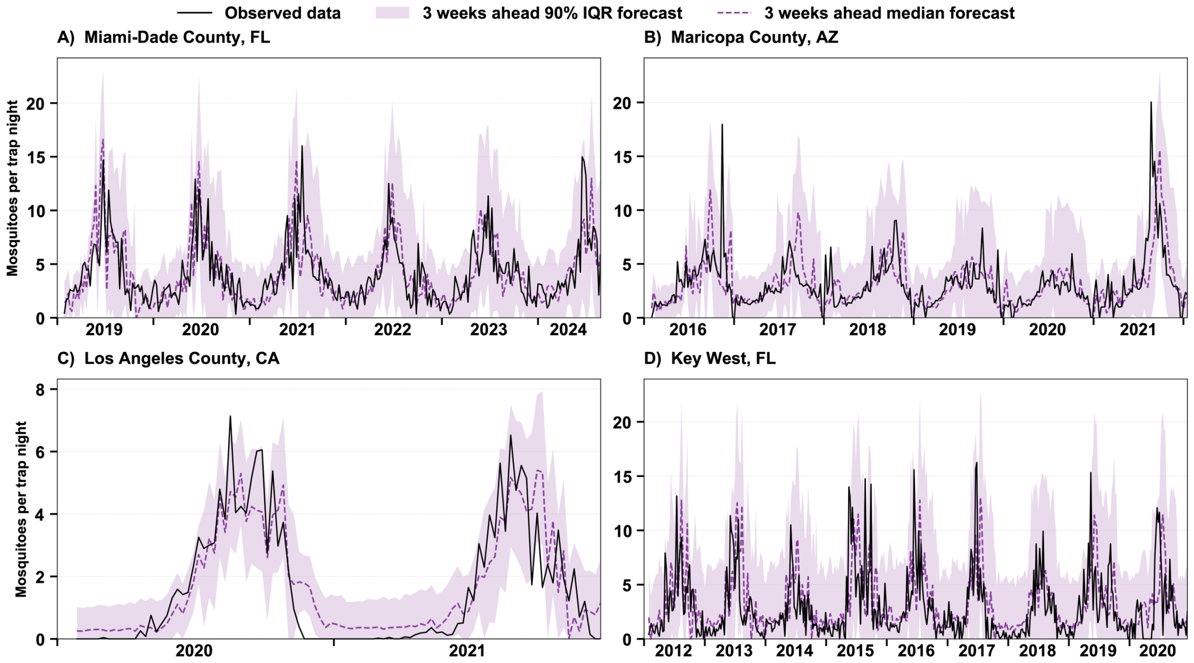

**Figure S11. Comparison between the observed *Ae. aegypti* relative abundance per trap night and 3-week-ahead forecasts (median and 90% prediction interval) for all forecast dates for the STL3 model. A.** Miami-Dade County, FL. **B.** Maricopa County, AZ. **C.** Los Angeles County, CA. **D.** Key West, FL.

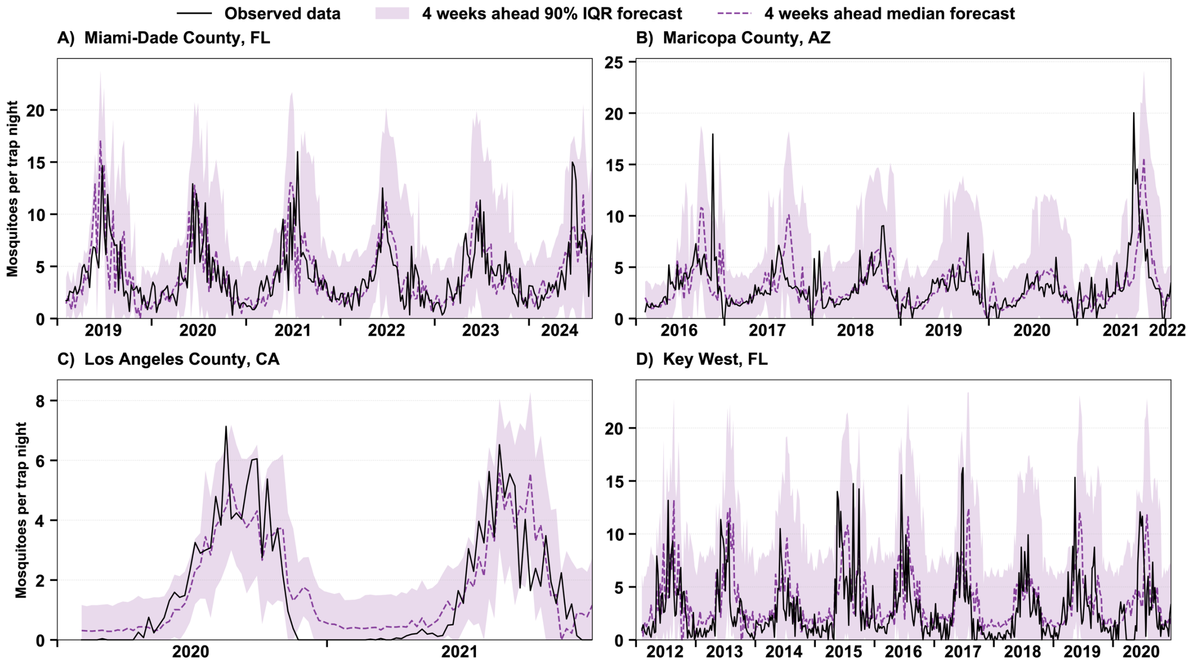

**Figure S12. Comparison between the observed *Ae. aegypti* relative abundance per trap night and 4-week-ahead forecasts (median and 90% prediction interval) for all forecast dates for the STL3 model. A.** Miami-Dade County, FL. **B.** Maricopa County, AZ. **C.** Los Angeles County, CA. **D.** Key West, FL.

### S2.1.4 STL4 model forecasts

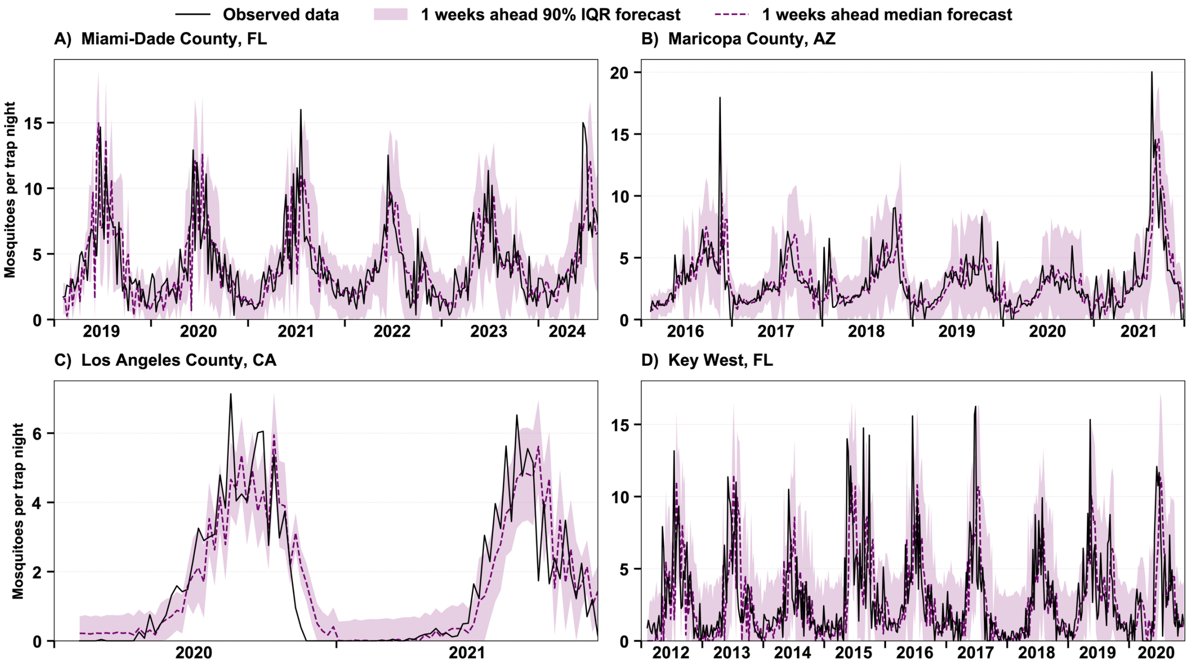

**Figure S13. Comparison between the observed *Ae. aegypti* relative abundance per trap night and 1-week-ahead forecasts (median and 90% prediction interval) for all forecast dates for the STL4 model. A.** Miami-Dade County, FL. **B.** Maricopa County, AZ. **C.** Los Angeles County, CA. **D.** Key West, FL.

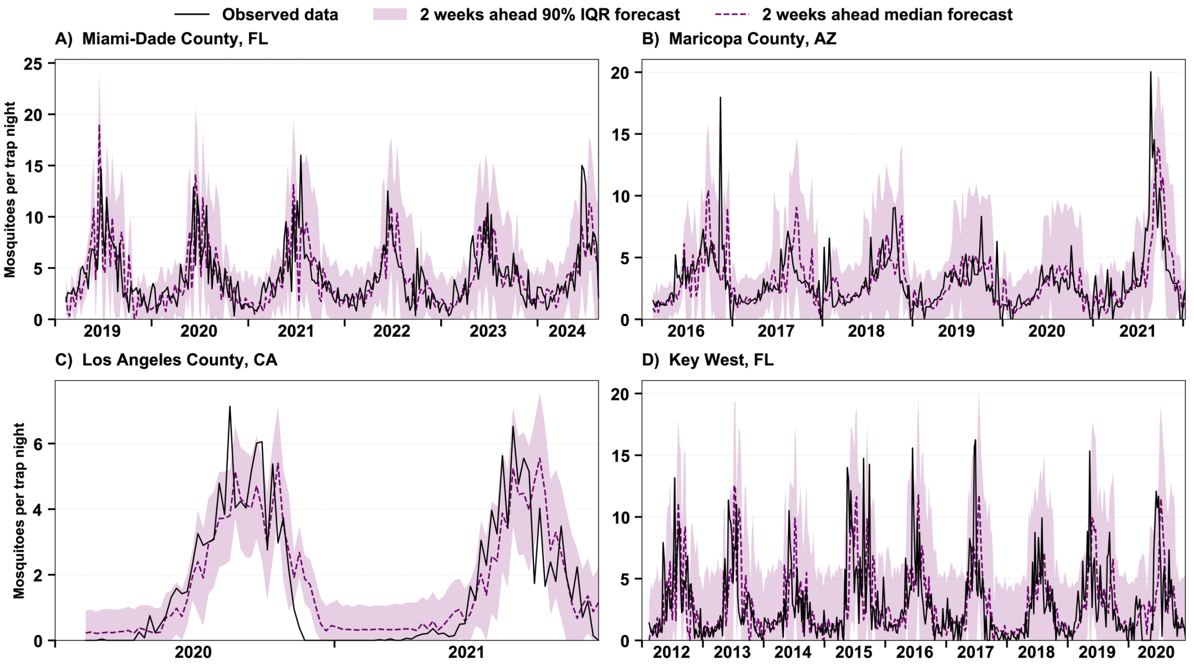

**Figure S14. Comparison between the observed *Ae. aegypti* relative abundance per trap night and 2-week-ahead forecasts (median and 90% prediction interval) for all forecast dates for the STL4 model. A.** Miami-Dade County, FL. **B.** Maricopa County, AZ. **C.** Los Angeles County, CA. **D.** Key West, FL.

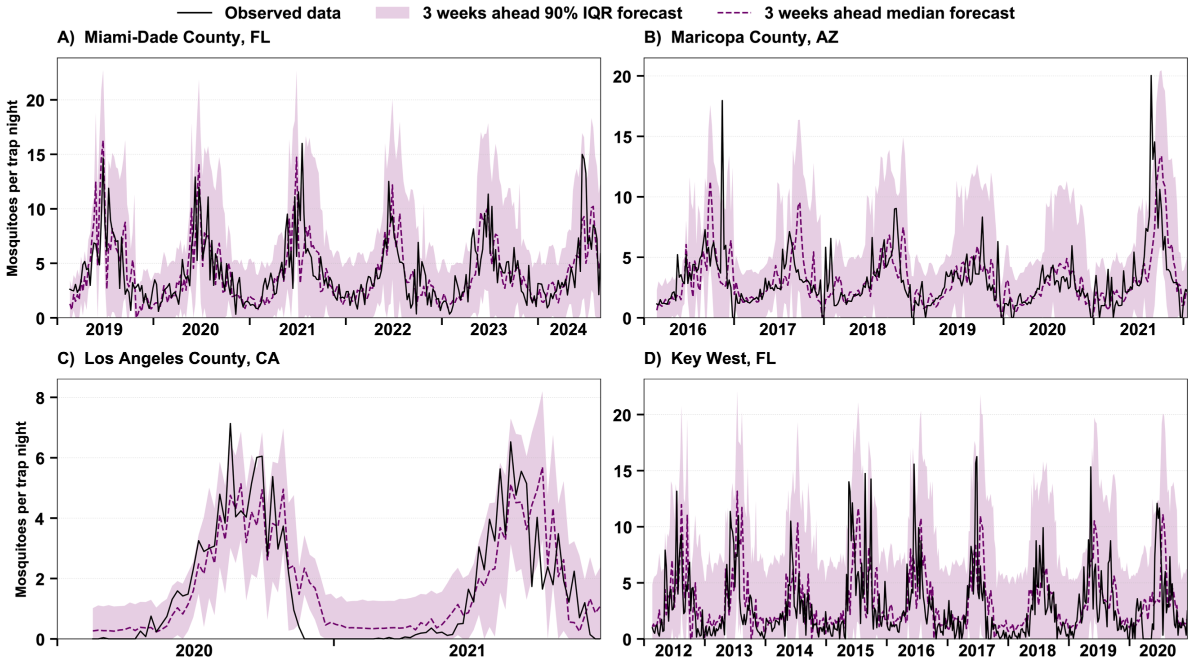

**Figure S15. Comparison between the observed *Ae. aegypti* relative abundance per trap night and 3-week-ahead forecasts (median and 90% prediction interval) for all forecast dates for the STL4 model. A.** Miami-Dade County, FL. **B.** Maricopa County, AZ. **C.** Los Angeles County, CA. **D.** Key West, FL.

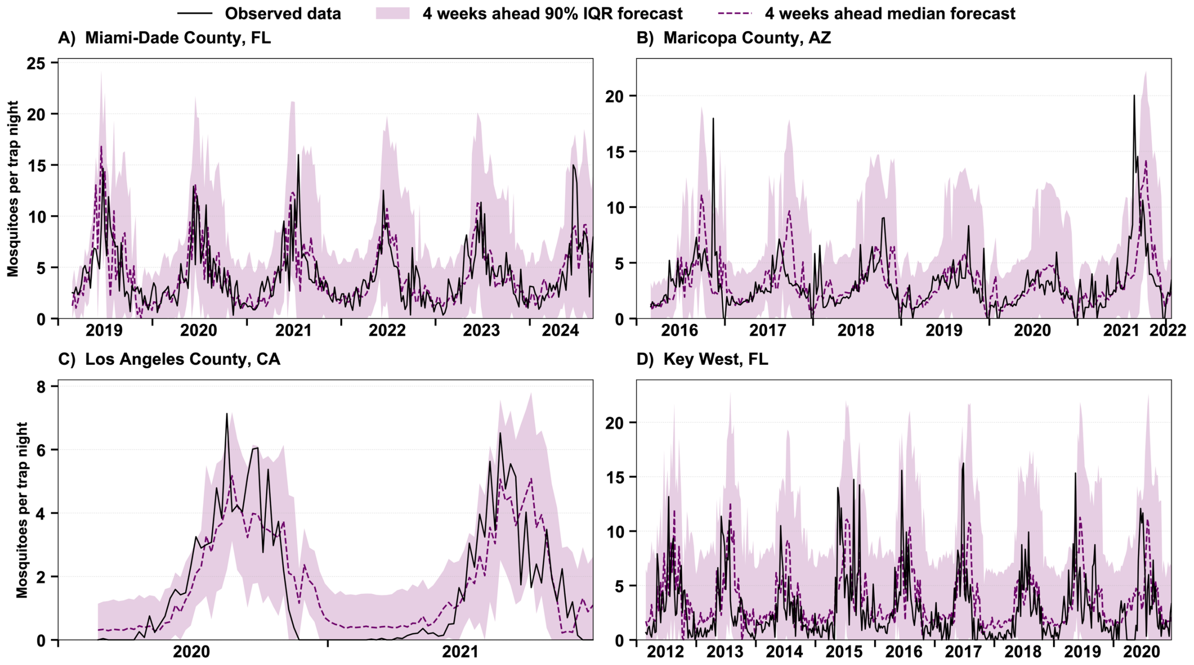

**Figure S16. Comparison between the observed *Ae. aegypti* relative abundance per trap night and 4-week-ahead forecasts (median and 90% prediction interval) for all forecast dates for the STL4 model. A.** Miami-Dade County, FL. **B.** Maricopa County, AZ. **C.** Los Angeles County, CA. **D.** Key West, FL.

### S2.1.5 SARIMAX model forecasts

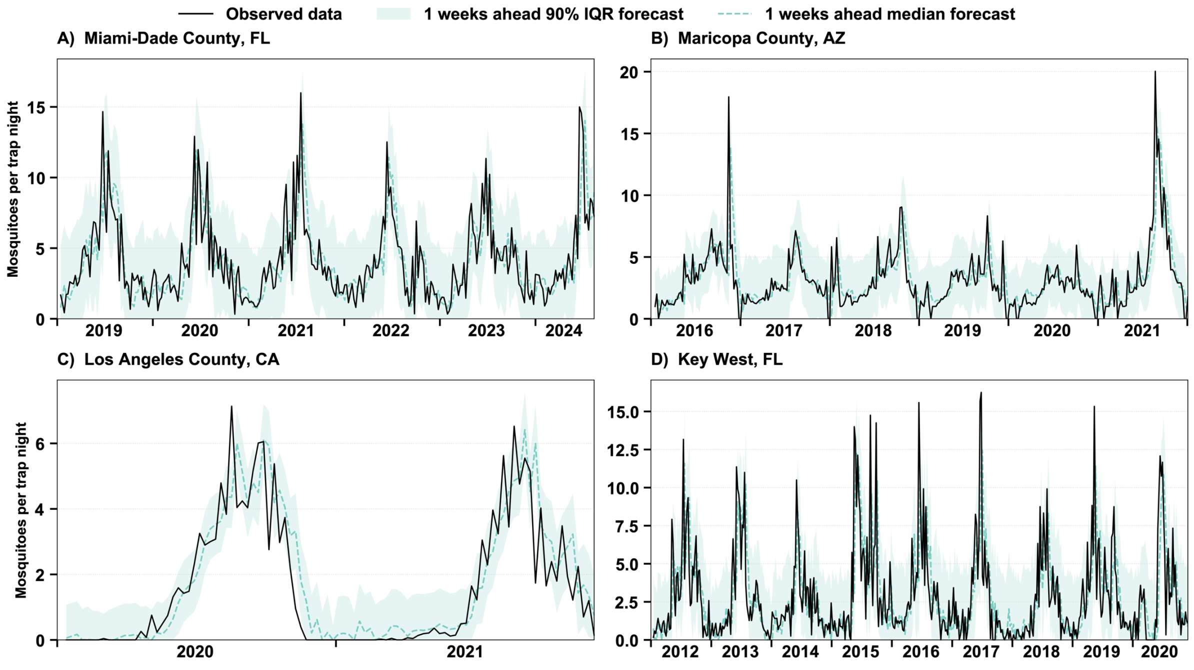

**Figure S17. Comparison between the observed *Ae. aegypti* relative abundance per trap night and 1-week-ahead forecasts (median and 90% prediction interval) for all forecast dates for the SARIMAX model. A.** Miami-Dade County, FL. **B.** Maricopa County, AZ. **C.** Los Angeles County, CA. **D.** Key West, FL.

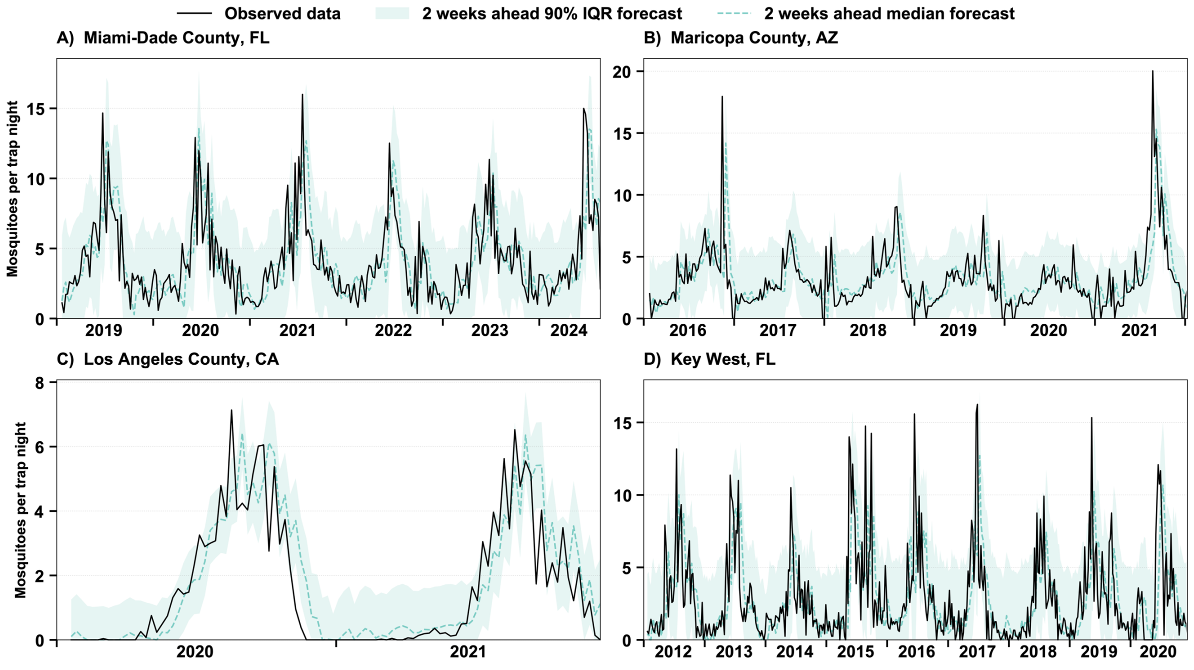

**Figure S18. Comparison between the observed *Ae. aegypti* relative abundance per trap night and 2-week-ahead forecasts (median and 90% prediction interval) for all forecast dates for the SARIMAX model. A.** Miami-Dade County, FL. **B.** Maricopa County, AZ. **C.** Los Angeles County, CA. **D.** Key West, FL.

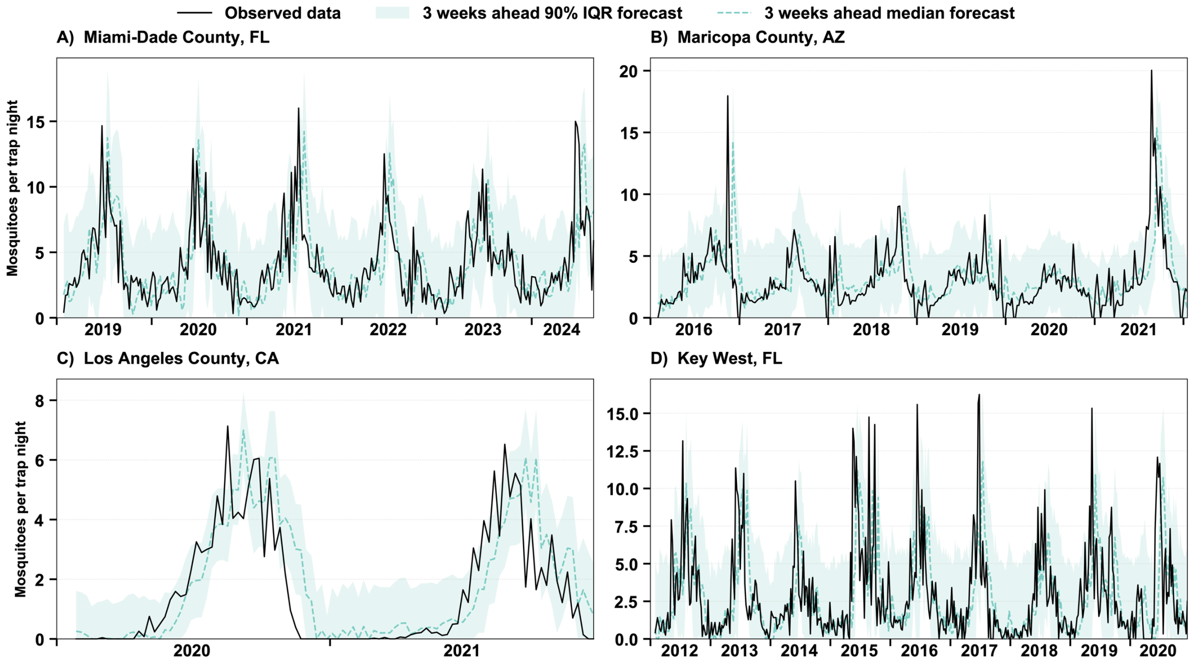

**Figure S19. Comparison between the observed *Ae. aegypti* relative abundance per trap night and 3-week-ahead forecasts (median and 90% prediction interval) for all forecast dates for the SARIMAX model. A.** Miami-Dade County, FL. **B.** Maricopa County, AZ. **C.** Los Angeles County, CA. **D.** Key West, FL.

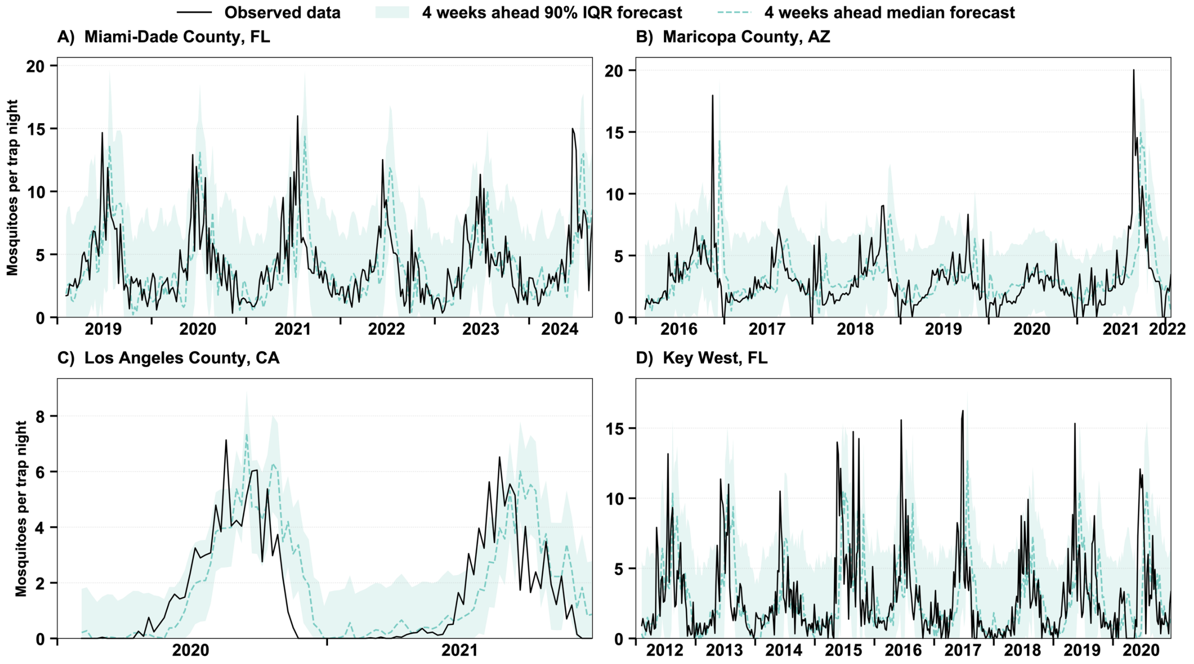

**Figure S20. Comparison between the observed *Ae. aegypti* relative abundance per trap night and 4-week-ahead forecasts (median and 90% prediction interval) for all forecast dates for the SARIMAX model. A.** Miami-Dade County, FL. **B.** Maricopa County, AZ. **C.** Los Angeles County, CA. **D.** Key West, FL.

### S2.1.6 UCM model forecasts

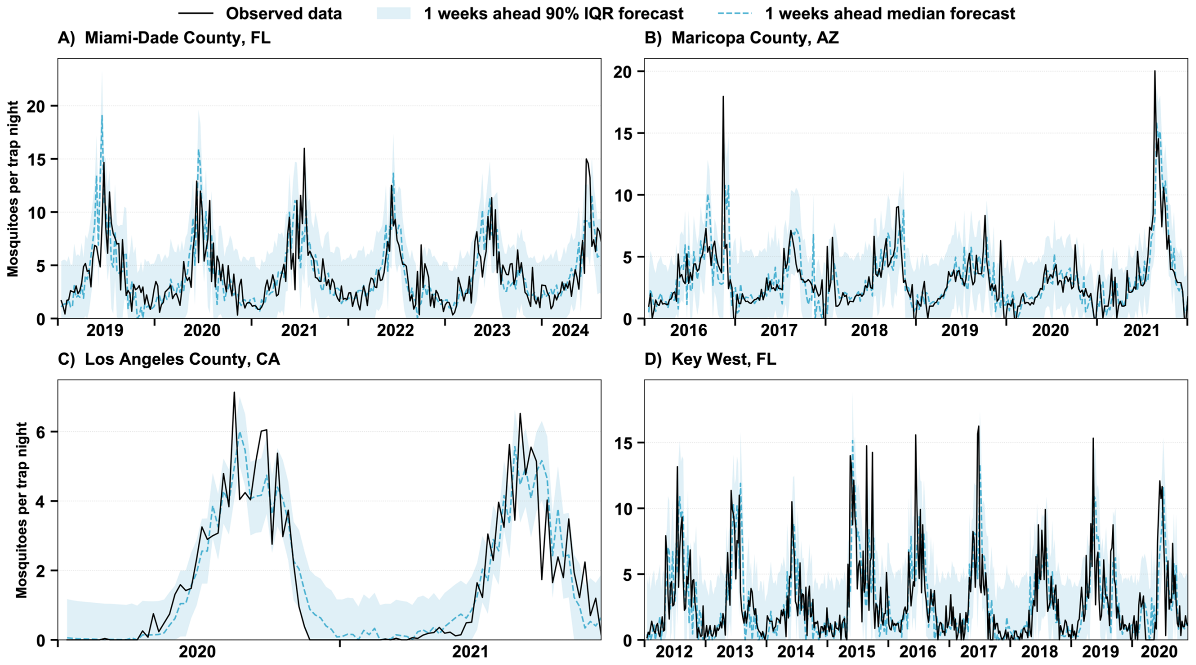

**Figure S21. Comparison between the observed *Ae. aegypti* relative abundance per trap night and 1-week-ahead forecasts (median and 90% prediction interval) for all forecast dates for the UCM model. A.** Miami-Dade County, FL. **B.** Maricopa County, AZ. **C.** Los Angeles County, CA. **D.** Key West, FL.

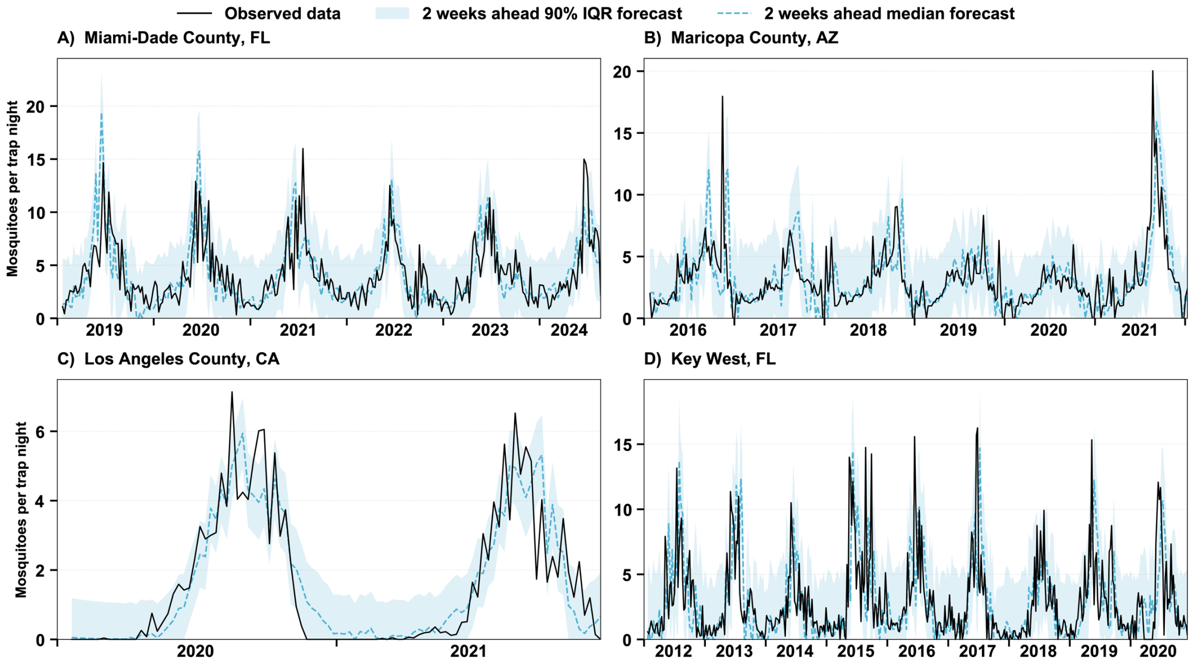

**Figure S22. Comparison between the observed *Ae. aegypti* relative abundance per trap night and 2-week-ahead forecasts (median and 90% prediction interval) for all forecast dates for the UCM model. A.** Miami-Dade County, FL. **B.** Maricopa County, AZ. **C.** Los Angeles County, CA. **D.** Key West, FL.

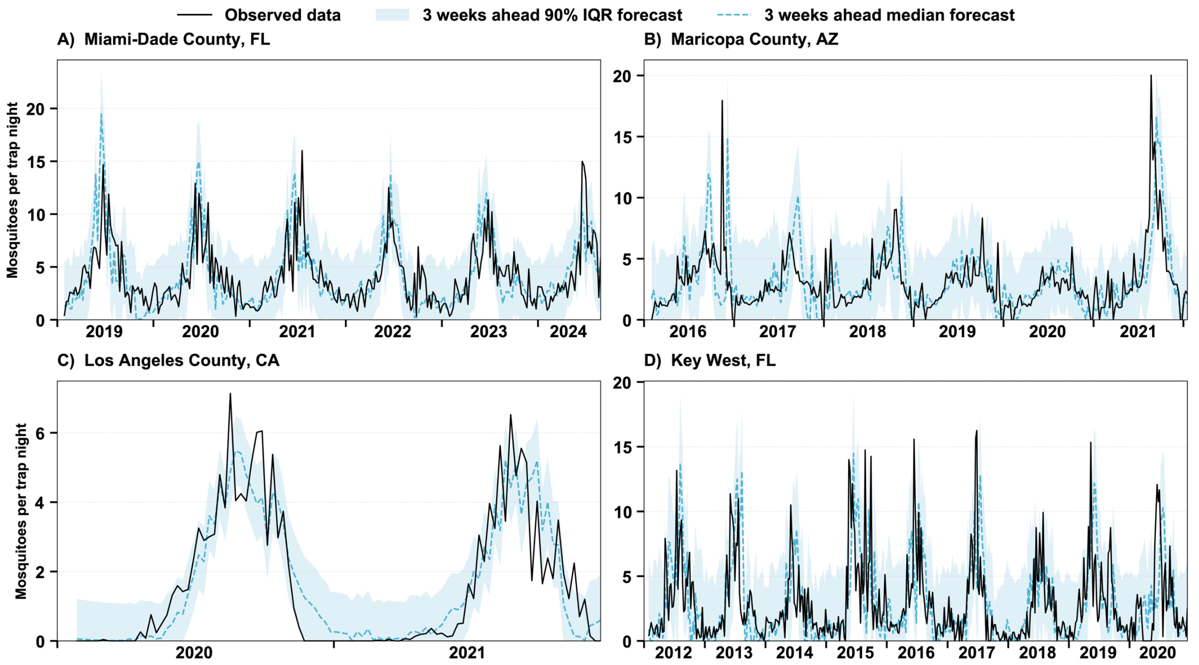

**Figure S23. Comparison between the observed *Ae. aegypti* relative abundance per trap night and 3-week-ahead forecasts (median and 90% prediction interval) for all forecast dates for the UCM model. A.** Miami-Dade County, FL. **B.** Maricopa County, AZ. **C.** Los Angeles County, CA. **D.** Key West, FL.

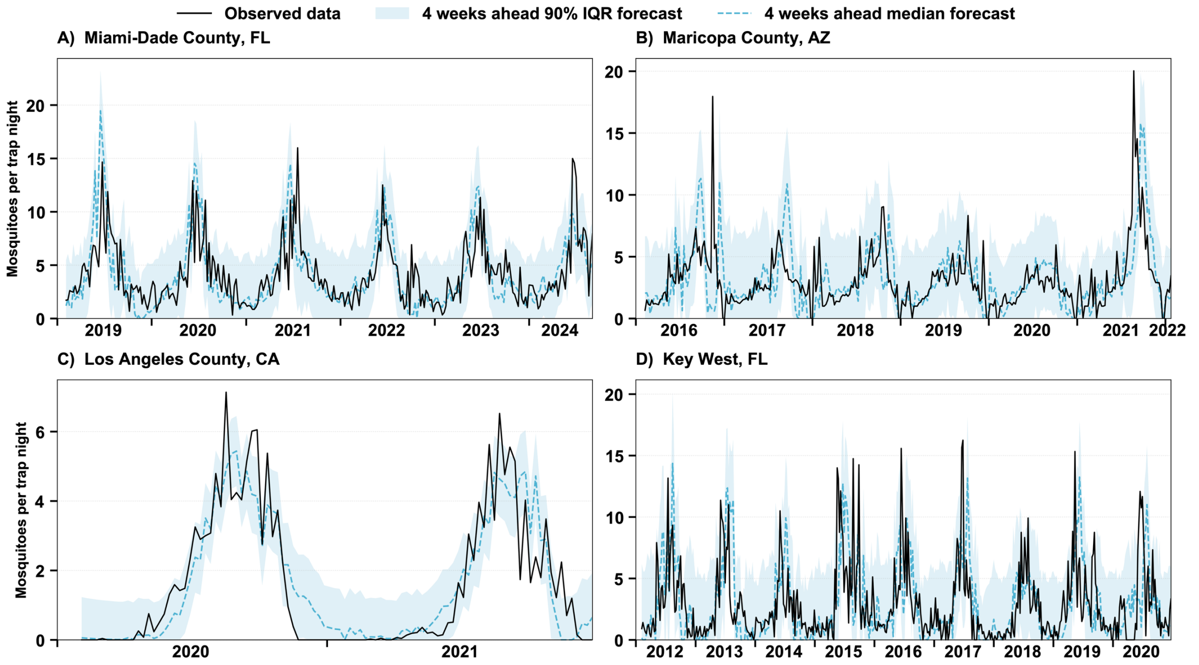

**Figure S24. Comparison between the observed *Ae. aegypti* relative abundance per trap night and 4-week-ahead forecasts (median and 90% prediction interval) for all forecast dates for the UCM model. A.** Miami-Dade County, FL. **B.** Maricopa County, AZ. **C.** Los Angeles County, CA. **D.** Key West, FL.

### S2.1.7 Ensemble model forecasts

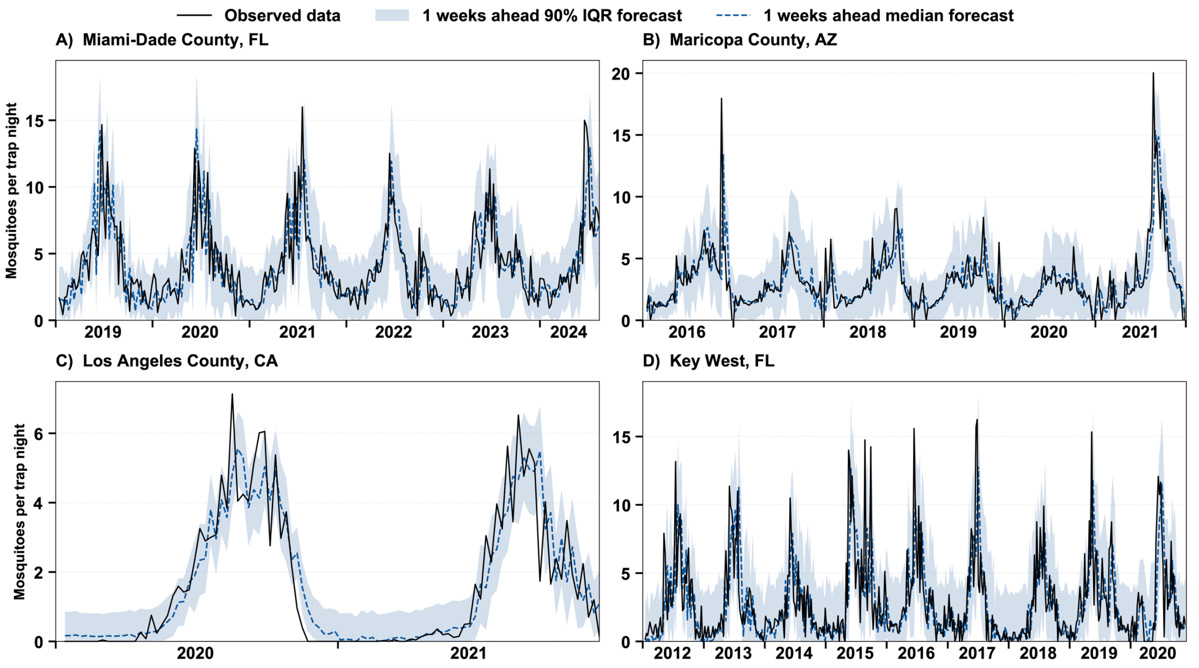

**Figure S25. Comparison between the observed *Ae. aegypti* relative abundance per trap night and 1-week-ahead forecasts (median and 90% prediction interval) for all forecast dates for the ensemble model. A.** Miami-Dade County, FL. **B.** Maricopa County, AZ. **C.** Los Angeles County, CA. **D.** Key West, FL.

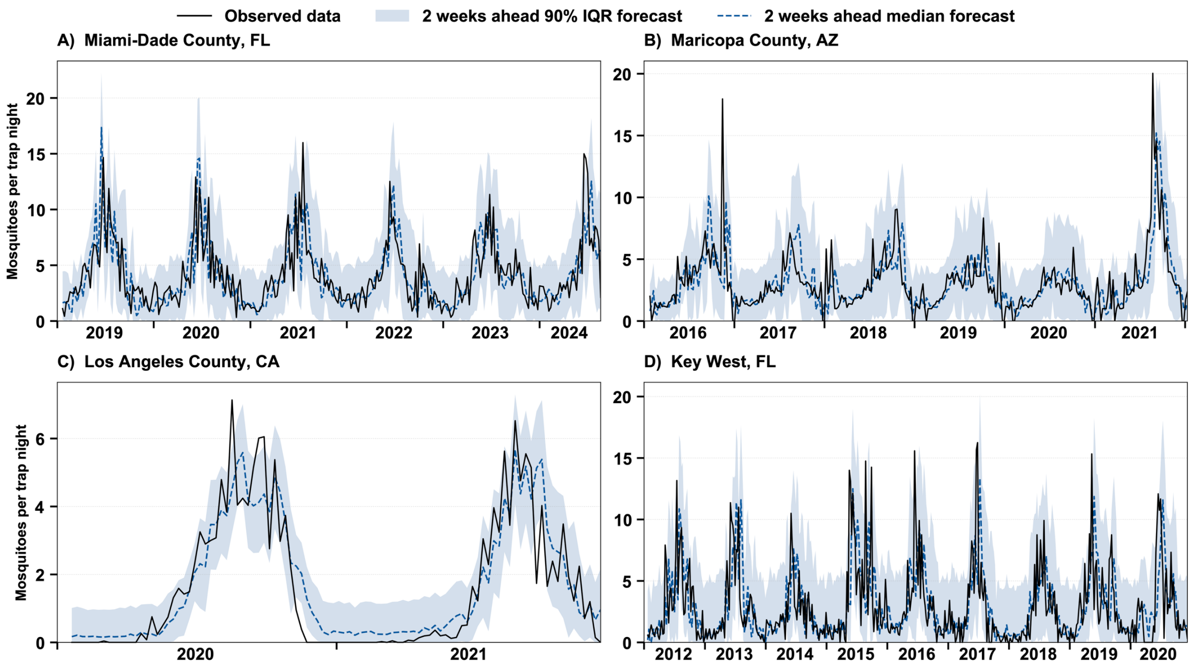

**Figure S26. Comparison between the observed *Ae. aegypti* relative abundance per trap night and 2-week-ahead forecasts (median and 90% prediction interval) for all forecast dates for the ensemble model. A.** Miami-Dade County, FL. **B.** Maricopa County, AZ. **C.** Los Angeles County, CA. **D.** Key West, FL.

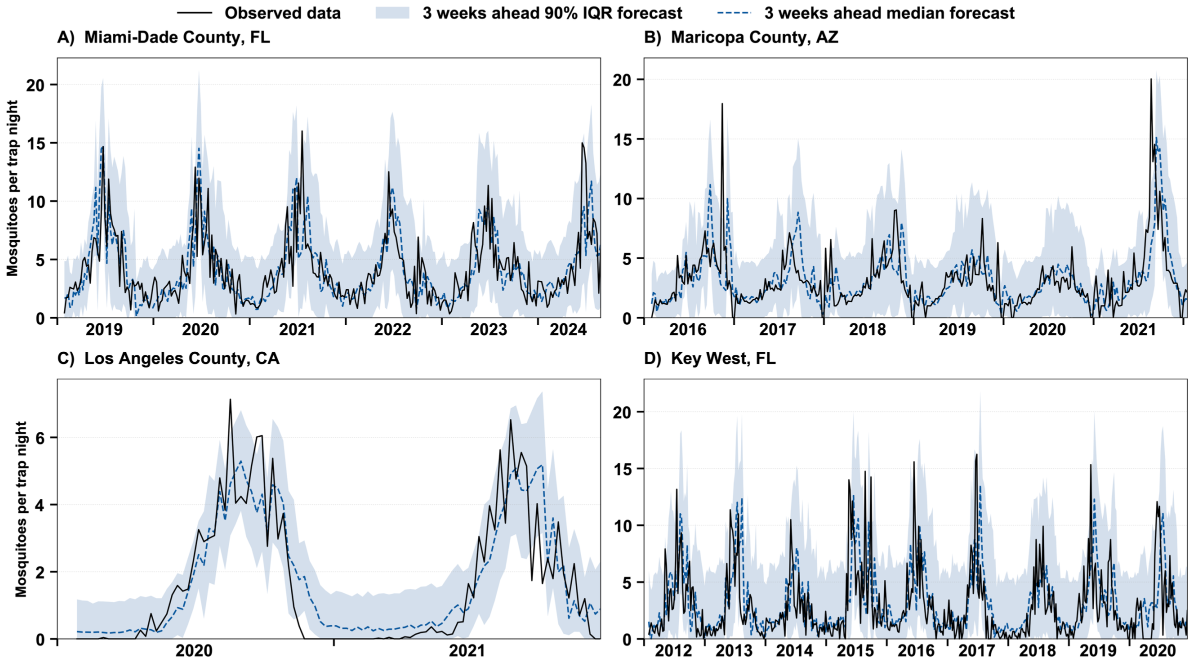

**Figure S27. Comparison between the observed *Ae. aegypti* relative abundance per trap night and 3-week-ahead forecasts (median and 90% prediction interval) for all forecast dates for the ensemble model. A.** Miami-Dade County, FL. **B.** Maricopa County, AZ. **C.** Los Angeles County, CA. **D.** Key West, FL.

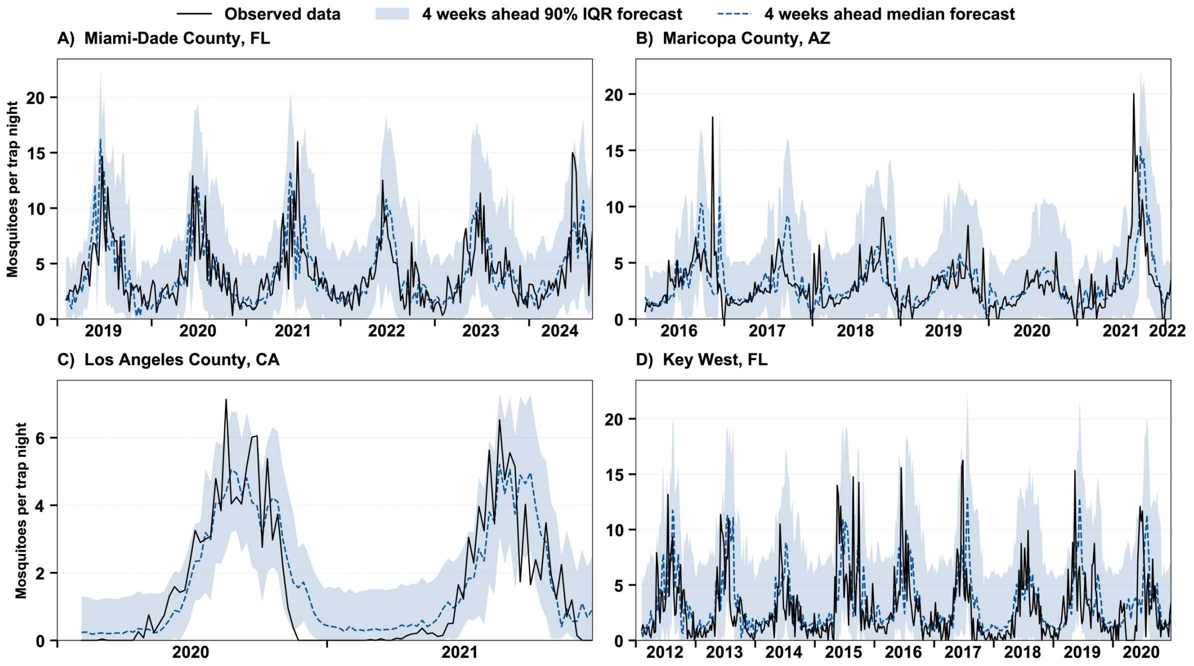

**Figure S28. Comparison between the observed *Ae. aegypti* relative abundance per trap night and 4-week-ahead forecasts (median and 90% prediction interval) for all forecast dates for the ensemble model. A.** Miami-Dade County, FL. **B.** Maricopa County, AZ. **C.** Los Angeles County, CA. **D.** Key West, FL.

### *S2.2 Estimates of performance metrics*

Estimates of the sMAPE, WIS, and 95% coverage for Key West, FL, Los Angeles County, CA, Maricopa County, AZ, and Miami-Dade County, FL, are shown in Tables S1-S4, respectively.

**Table S1. Evaluation scores of all models for Key West for all week-ahead horizons**

| Model | sMAPE | | | | WIS | | | | Coverage 95% | | | |
|---|---|---|---|---|---|---|---|---|---|---|---|---|
| | 1 | 2 | 3 | 4 | 1 | 2 | 3 | 4 | 1 | 2 | 3 | 4 |
| Naïve | 69.57 | 77.23 | 78.99 | 82.26 | 4.79 | 5.66 | 5.96 | 6.35 | 88.7 | 91.1 | 92.4 | 94.4 |
| Seasonal_1 | 80.31 | 71.26 | 71.10 | 74.20 | 4.67 | 5.31 | 5.61 | 6.07 | 95.0 | 96.8 | 98.5 | 98.9 |
| Seasonal_2 | 77.97 | 70.00 | 71.48 | 73.59 | 4.67 | 5.20 | 5.50 | 6.00 | 94.6 | 96.8 | 98.7 | 98.9 |
| Seasonal_3 | 76.47 | 69.81 | 70.86 | 73.90 | 4.72 | 5.21 | 5.55 | 5.99 | 94.4 | 97.4 | 98.7 | 99.1 |
| Seasonal_4 | 77.54 | 70.59 | 70.36 | 75.0 | 4.85 | 5.35 | 5.62 | 6.0 | 94.4 | 97.8 | 98.7 | 99.1 |
| SARIMAX | 69.02 | 69.70 | 73.89 | 76.04 | 4.51 | 5.11 | 5.49 | 5.80 | 93.3 | 92.2 | 93.3 | 93.8 |
| UCM | 72.49 | 78.42 | 83.08 | 86.72 | 4.74 | 5.28 | 5.57 | 5.81 | 93.8 | 93.1 | 92.9 | 92.0 |
| Ensemble | 68.22 | 65.87 | 67.30 | 69.44 | 4.40 | 4.97 | 5.25 | 5.61 | 94.6 | 96.8 | 97.4 | 98.3 |

**Table S2. Evaluation scores of all models for Los Angeles County for all week-ahead horizons**

| Model | sMAPE | | | | WIS | | | | Coverage 95% | | | |
|---|---|---|---|---|---|---|---|---|---|---|---|---|
| | 1 | 2 | 3 | 4 | 1 | 2 | 3 | 4 | 1 | 2 | 3 | 4 |
| Naïve | 71.31 | 77.42 | 84.54 | 97.37 | 2.14 | 1.85 | 2.66 | 2.84 | 82.0 | 91.0 | 87.0 | 93.0 |
| Seasonal_1 | 87.59 | 93.55 | 105.36 | 98.03 | 1.99 | 1.49 | 1.94 | 1.84 | 79.0 | 94.0 | 96.0 | 95.0 |
| Seasonal_2 | 83.31 | 95.58 | 98.63 | 99.12 | 1.50 | 1.64 | 1.77 | 1.81 | 89.0 | 94.0 | 95.0 | 96.0 |
| Seasonal_3 | 85.29 | 95.49 | 99.84 | 100.93 | 1.72 | 1.74 | 1.86 | 1.96 | 83.0 | 90.0 | 95.0 | 95.0 |
| Seasonal_4 | 81.52 | 93.22 | 96.89 | 98.98 | 1.83 | 1.91 | 2.03 | 2.16 | 82.47 | 89.69 | 93.81 | 94.84 |
| ARIMA | 78.88 | 78.74 | 88.47 | 93.74 | 1.77 | 1.86 | 2.47 | 2.72 | 86.0 | 89.0 | 86.0 | 84.0 |
| UCM | 90.77 | 93.90 | 99.89 | 100.41 | 1.65 | 1.67 | 1.84 | 1.81 | 89.0 | 87.0 | 86.0 | 87.0 |
| Ensemble | 88.37 | 93.25 | 97.0 | 98.33 | 1.58 | 1.57 | 1.8 | 1.86 | 86.0 | 92.0 | 92.0 | 93.0 |

**Table S3. Evaluation scores of all models for Maricopa County for all week-ahead horizons.**

| Model | sMAPE | | | | WIS | | | | Coverage 95% | | | |
|---|---|---|---|---|---|---|---|---|---|---|---|---|
| | 1 | 2 | 3 | 4 | 1 | 2 | 3 | 4 | 1 | 2 | 3 | 4 |
| Naïve | 41.09 | 51.31 | 55.28 | 58.19 | 3.36 | 4.09 | 4.48 | 4.92 | 92.63 | 95.83 | 96.47 | 97.44 |
| Seasonal_1 | 38.16 | 48.51 | 43.40 | 46.74 | 3.52 | 4.12 | 4.29 | 4.71 | 91.35 | 92.95 | 95.51 | 96.15 |
| Seasonal_2 | 41.36 | 44.17 | 43.64 | 47.78 | 3.30 | 3.89 | 4.20 | 4.53 | 91.35 | 93.91 | 96.80 | 97.44 |
| Seasonal_3 | 40.30 | 44.56 | 44.69 | 46.09 | 3.31 | 3.95 | 4.21 | 4.49 | 90.06 | 94.87 | 97.44 | 97.12 |
| Seasonal_4 | 39.94 | 45.22 | 44.76 | 45.93 | 3.40 | 4.00 | 4.21 | 4.50 | 92.88 | 94.82 | 97.09 | 97.74 |
| ARIMA | 37.94 | 42.47 | 46.17 | 48.64 | 3.28 | 3.85 | 4.28 | 4.63 | 94.55 | 94.87 | 94.55 | 92.95 |
| UCM | 45.45 | 49.83 | 51.74 | 54.25 | 3.66 | 4.19 | 4.63 | 4.93 | 94.87 | 93.91 | 94.55 | 93.27 |
| Ensemble | 38.10 | 43.34 | 45.15 | 46.88 | 3.20 | 3.75 | 4.09 | 4.40 | 92.63 | 94.55 | 96.47 | 97.12 |

**Table S4. Evaluation scores of all models for Miami-Dade County for all week-ahead horizons**

| Model | sMAPE | | | | WIS | | | | Coverage 95% | | | |
|---|---|---|---|---|---|---|---|---|---|---|---|---|
| | 1 | 2 | 3 | 4 | 1 | 2 | 3 | 4 | 1 | 2 | 3 | 4 |
| Naïve | 39.31 | 44.16 | 48.54 | 50.97 | 4.71 | 5.05 | 5.90 | 6.09 | 91.78 | 95.55 | 94.52 | 97.60 |
| Seasonal_1 | 44.46 | 46.13 | 46.19 | 45.45 | 4.95 | 4.96 | 5.18 | 5.25 | 86.64 | 95.21 | 96.92 | 97.60 |
| Seasonal_2 | 39.97 | 42.89 | 42.48 | 42.85 | 4.47 | 4.81 | 4.80 | 5.10 | 89.04 | 95.89 | 97.95 | 98.29 |
| Seasonal_3 | 39.86 | 42.05 | 41.38 | 41.65 | 4.56 | 4.74 | 4.79 | 5.05 | 89.38 | 95.21 | 98.63 | 98.63 |
| Seasonal_4 | 39.79 | 42.04 | 41.09 | 40.50 | 4.54 | 4.80 | 4.77 | 5.01 | 89.97 | 95.16 | 98.27 | 98.62 |
| ARIMA | 35.42 | 39.23 | 44.33 | 47.33 | 4.17 | 4.58 | 5.20 | 5.54 | 94.52 | 94.18 | 93.84 | 95.55 |
| UCM | 39.67 | 41.94 | 44.51 | 45.65 | 4.69 | 4.85 | 5.14 | 5.25 | 93.84 | 94.52 | 93.49 | 93.49 |
| Ensemble | 36.35 | 38.32 | 39.47 | 40.14 | 4.17 | 4.38 | 4.52 | 4.71 | 93.84 | 97.26 | 98.97 | 99.32 |
